# Full-length structure of the anti-viral and pro-tumor DNA deaminase APOBEC3B

**DOI:** 10.64898/2026.06.18.733170

**Authors:** Ryan H. Abdella, Christopher A. Belica, Yanjun Chen, William L. Brown, Michael A. Carpenter, Mahmoud A. Ibrahim, Bárbara de la Peña Avalos, Christopher D. Mullally, Allen J. York, Reuben S. Harris, Hideki Aihara

## Abstract

Human APOBEC3B (A3B) restricts virus infections by catalyzing the deamination of cytosines to uracils in single-stranded DNA. A3B also contributes to mutagenesis and genome instability in cancer cells, driving tumor evolution and detrimental outcomes including therapy resistance and metastasis. A3B comprises tandem globular deaminase domains, with a multifunctional amino-terminal domain (NTD) and a catalytically active carboxy-terminal domain (CTD). Although individual domain structures have been studied, the structure of full-length A3B has remained elusive. Here, we report the cryoEM structure of wildtype A3B in complex with the natural antagonist BORF2 (the large subunit of the Epstein-Barr virus ribonucleotide reductase). The two domains of A3B bridge a novel BORF2 dimer interface, showing a unique domain positioning that distinguishes A3B from the related dual-domain retrovirus restriction factor APOBEC3G (A3G). Mutational analyses suggest that the unique NTD-CTD interaction regulates A3B deaminase activity. The BORF2 dimerization interface is stabilized by primary interactions with A3B-CTD and secondary contacts with A3B-NTD, as well as by A3B CTD-CTD dimerization. This matrix of interactions supports a molecular mechanism for A3B neutralization in which BORF2 binding leads to deaminase sequestration in large aggregates. The full-length wildtype A3B structure also provides a platform for future anti-viral and anti-cancer drug development efforts.

## Introduction

The human APOBEC3 (A3) family of single-stranded (ss)DNA cytosine deaminases (A3A, A3B, A3C, A3D, A3F, A3G, and A3H) are innate immune enzymes that provide an overlapping defense against a wide range of DNA viruses, retroviruses, and retrotransposons^1–5^. Specific examples include HIV-1, hepatitis B virus, Epstein-Barr virus, and human papillomavirus. Virus restriction is mediated in part by catalyzing hydrolytic deamination of cytosines to uracils (C-to-U) and thereby causing hypermutation of viral genomes. Each A3 protein is comprised of a single zinc (Zn)-bound deaminase domain, as for A3A, A3C, and A3H, or two tandemly arranged deaminase-fold domains, as for A3B, A3D, A3F, and A3G^6,7^. In the human dual-domain A3 proteins, the N-terminal domain (NTD) is catalytically inactive and provides regulatory functions, whereas the C-terminal domain (CTD) catalyzes C-to-U deamination. The catalytic domains of A3s engage nucleic acid substrate through three flexible loops surrounding the active site (loops 1, 3, and 7), which govern preferences for distinct sequence motifs in ssDNA^8–10^.

A3B is the only nuclear-localized family member, with the NTD mediating a non-canonical mechanism of nuclear import^11–14^. A3B has been strongly implicated in restricting herpesvirus replication, which occurs in the nucleus of infected cells^15–17^. To counteract the antiviral activity of A3B and preserve viral genomic DNA integrity, herpesviruses have evolved a unique mechanism in which the large subunit of the viral ribonucleotide reductase (RNRα) binds tightly to the active site of A3Bctd to block DNA deamination activity^15,18^. At the same time, the RNRα triggers the sequestration of A3B into large cytoplasmic complexes to further prevent access to viral DNA in the nucleus^15–17^. Structural studies have shown that the RNRα protein of the gamma-herpesvirus EBV (BORF2) binds to A3B CTD loops 1 and 7, which prevents ssDNA from accessing the active site^18^. However, how EBV BORF2 binds to and effectively neutralizes the full-length A3B protein, which contains the NTD responsible for nuclear import, and the precise nature of the higher-order A3B-BORF2 aggregates formed in the cytoplasm remain open questions.

A3B is also dysregulated in a range of different cancers, causing genomic DNA mutations and chromosomal instability associated with detrimental outcomes, including therapy resistance and metastasis^19–25^. A3B signature mutations, C-to-T and C-to-G in 5’-TC dinucleotides, are evident in nearly 70% of human cancer types, including those of breast, bladder, cervix, head/neck, and lung^20,26,27^. Moreover, the mutagenic activity of A3B is exacerbated by chemical carcinogens such as those in tobacco smoke^28^. A3B is therefore a prime target for therapeutic inhibition, which is predicted to slow tumor evolvability and prevent mortality. However, despite nearly two decades of research, the structure of full-length A3B remains unknown, limiting our understanding of how the two domains coordinate intracellularly and the possibility of structure-based inhibitor design. Here, we present the cryoEM structure of full-length, wildtype human A3B (A3Bwt) in complex with EBV BORF2, which provides the first view of A3B complete with both deaminase-fold domains and also informs on how EBV sequesters A3B to circumvent restriction.

## Results

### Purification and characterization of A3Bwt

Attempts to characterize the structure of A3Bwt have been unsuccessful due to low solubility and a high propensity to aggregate, which is largely caused by the NTD^29,30^. A3Bwt was expressed in Expi293T cells, and buffer conditions were evaluated systematically to improve solubility for structural studies. Adding 50 mM L-arginine and adjusting buffer pH to 9.0 increased the amount of A3Bwt in clarified lysates (**Extended Data Figure 1a,b**). A3Bwt also required abnormally high imidazole concentrations to elute from Ni-NTA resin and it eluted slowly (**Extended Data Figure 1c**), suggesting that some of the purified protein may be oligomeric and agreeing with work showing that A3Bwt can form high molecular weight aggregates^29–31^. Size exclusion chromatography confirmed that A3Bwt purifies as two distinct species: a high molecular weight peak with a high A260/A280 ratio and co-eluting proteins, and a monomeric peak with a lower A260/A280 ratio (**Figure 1a**). All subsequent experiments were performed using the monomeric A3Bwt peak, except for cryoEM studies, which required an entire Ni-NTA eluate as input. The purified A3Bwt exhibited robust DNA deaminase activity in the real-time APOBEC-mediated DNA deamination (RADD) assay^32,33^ (**Figure 1b**). Moreover, BORF2 dose-responsively inhibits the DNA deaminase activity of A3Bwt (**Figure 1c**), analogous to prior observations with BORF2 and A3Bctd alone^15,18^. These experiments show that the A3Bwt protein is active and capable of interaction with a biologically relevant partner.

**Figure 1:**
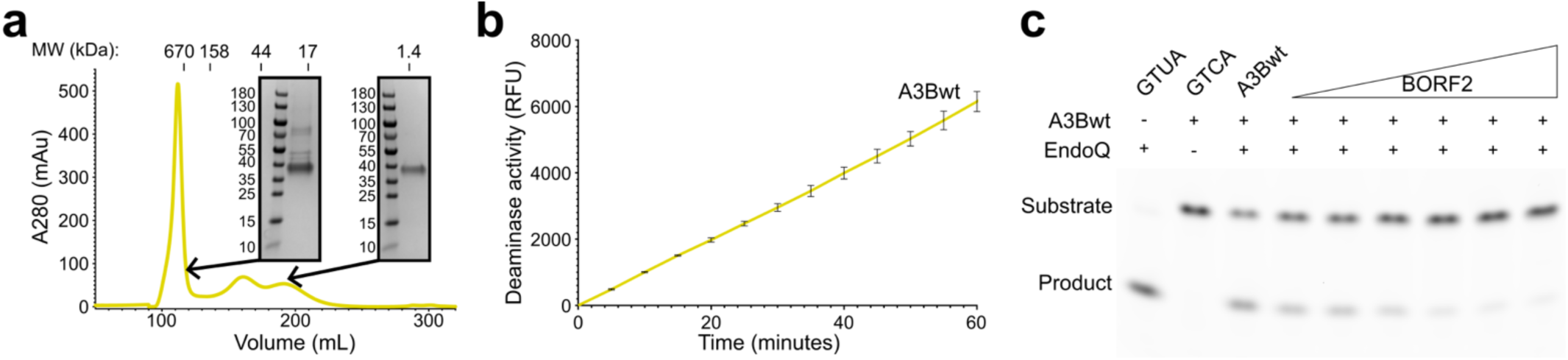
Wildtype A3B purification, deaminase activity, and inhibition by EBV BORF2. **a**, Size-exclusion chromatogram for A3Bwt. The Coomassie-stained SDS-PAGE images show contaminants in the early-eluting peak and a more homogenous late-eluting peak. The intermediate peak may represent a dimer. **b**, DNA deaminase activity of 180 nM A3Bwt in the RADD assay (n=3 experiments with mean+/-SD shown). **c**, Dose-responsive inhibition of A3Bwt by BORF2 in a gel-based ssDNA deamination assay (180 nM A3Bwt and 0, 25, 50, 100, 200, 400, and 800 nM BORF2). The control reaction with full cleavage (GTUA) shows that EndoQ is not rate-limiting, and the reaction with no cleavage (GTCA) shows that EndoQ is essential.

### A3Bwt-BORF2 cryoEM complexes

For structural studies, A3Bwt was incubated with EBV BORF2, and the resulting complexes were purified by size exclusion chromatography (**Figure 2a,b**). The A3Bwt-BORF2 complex eluted at a very high molecular weight (ca. 700 kDa). Therefore, all fractions that contained both proteins were pooled, concentrated to 0.5 mg/mL, and used to prepare grids for cryoEM. Systematic data processing yielded structures of a tetrameric A3Bwt-BORF2 complex with two copies of each protein at 2.67 Å resolution (**Figure 2c**, **Table 1**, and **Extended Data Figure 2**). The complex was symmetry expanded and subjected to 3D classification with 2 classes and a mask around A3Bwt. Particles from the class showing better density for A3Bwt were selected. All subsequent steps were performed on the symmetry expanded particle stack. Focused refinements were performed on A3Bwt and BORF2, which resulted in a 2.8 Å map of A3Bwt that showed improved density for A3Bntd and a map at 2.4 Å resolution of BORF2. A focused refinement on the A3Bwt-BORF2 heterodimer resulted in a map at 2.6 Å (**Table 1** and **Extended Data Figure 2**).

**Figure 2:**
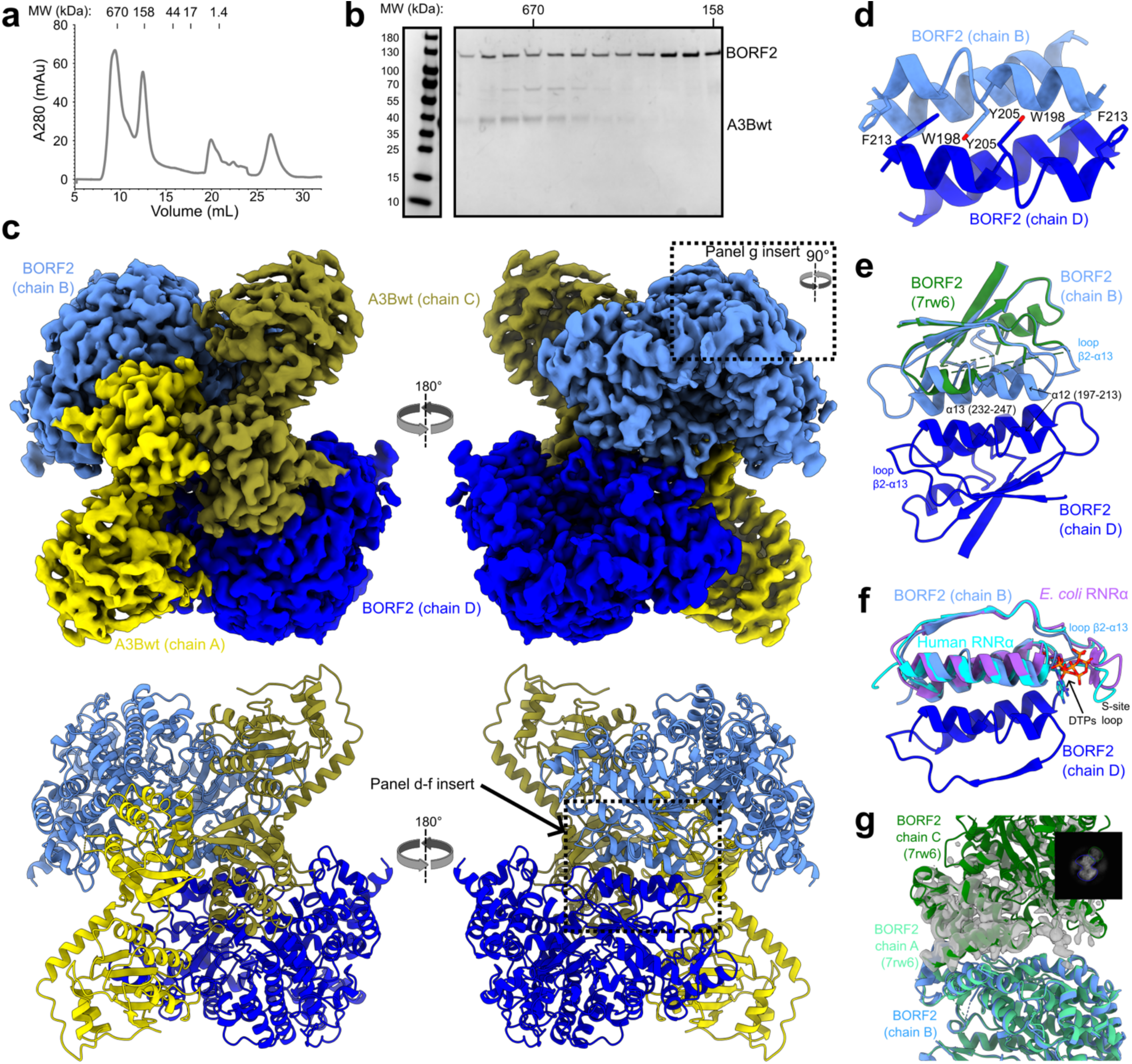
Overall structure of the A3Bwt-BORF2 complex. **a**, Size-exclusion chromatogram of the A3Bwt-BORF2 complex. **b**, Coomassie-stained SDS-PAGE of relevant fractions from panel a. **c**, CryoEM map (top) and ribbon model (bottom) of the heterotetrameric A3Bwt-BORF2 complex (two views, rotated 180°). A3Bwt chains A and C are shown in yellow and gold, and BORF2 chains B and D are shown in light and dark blue. Dashed boxes indicate regions expanded in panels d-g. **d**, Ribbon diagram of the canonical BORF2 dimer interface formed by α-helices 12 and 13. Aromatic side chains involved in π-π stacking interactions are indicated. **e**, An expanded ribbon view of the canonical BORF2 dimer interface overlaid with the same portion of BORF2 from the A3Bctd-BORF2 complex^18^ (pdb: 7rw6). **f**, A ribbon diagram of the canonical BORF2 dimer interface aligned with the corresponding region of the human RNRα subunit^61^ (pdb: 6aui, cyan) and *E. coli* RNRα subunit^62^ (pdb: 9db2, magenta) with DTP in the S-site. **g**, Ribbon schematic of the noncanonical BORF2 dimer interface. Chain A of BORF2 (light green) from the A3Bctd-BORF2 complex^18^ (pdb: 7rw6) was aligned to chain B of BORF2 from the A3Bwt-BORF2 complex (light blue). Chain C of BORF2 from the A3Bctd-BORF2 complex fits well into the segmented cryoEM density (transparent gray) not accounted for in the A3Bwt-BORF2 model. Insert: Representative 2D class from the A3Bwt-BORF2 dataset with the canonical BORF2 dimer outlined in blue and additional density for an extra copy of BORF2 bound via the non-canonical dimer interface in green.

**Table 1:**
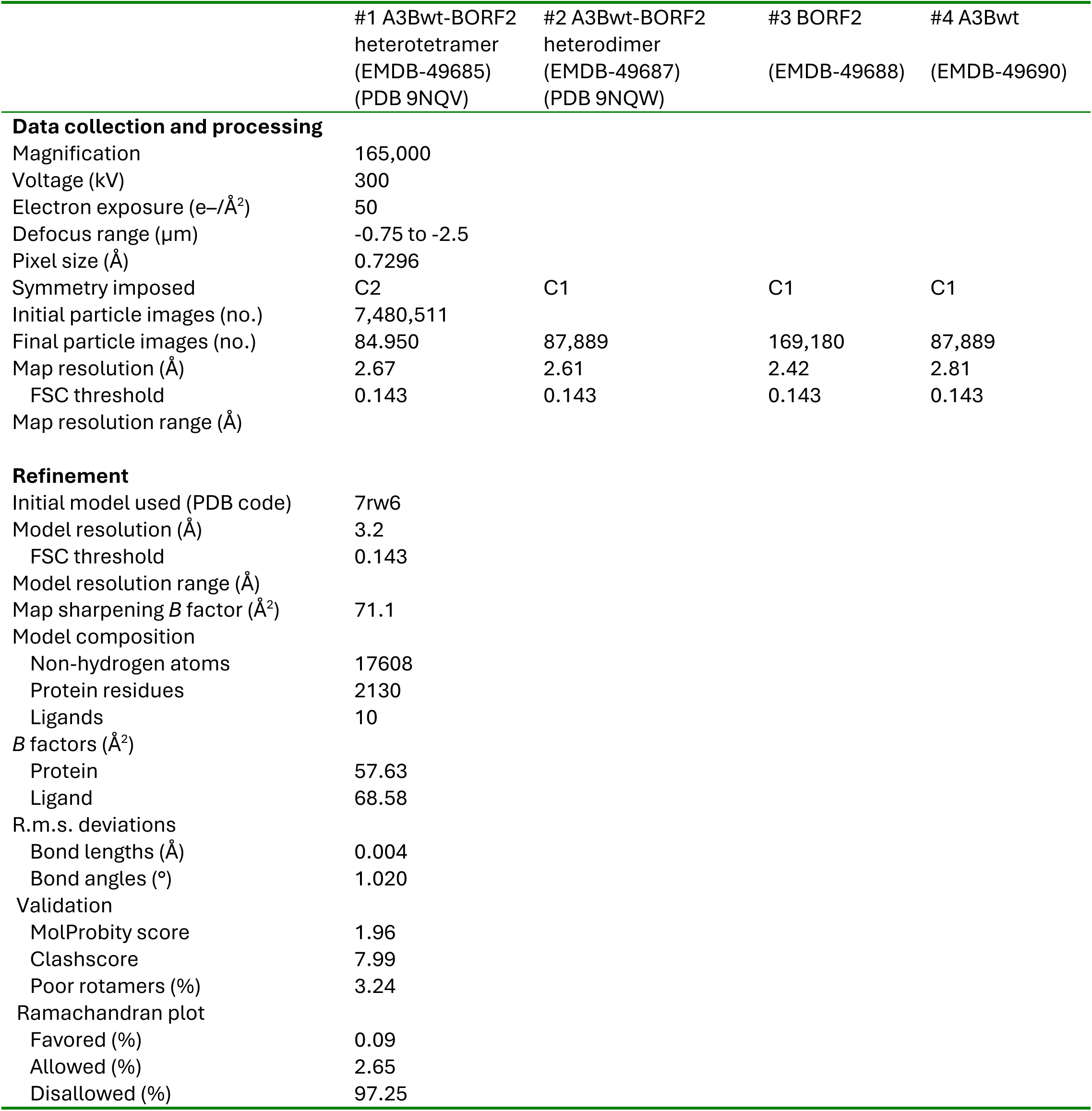
Cryo-EM data collection, refinement and validation statistics.

|  | #1 A3Bwt-BORF2<br>heterotetramer<br>(EMDB-49685)<br>(PDB 9NQV) | #2 A3Bwt-BORF2<br>heterodimer<br>(EMDB-49687)<br>(PDB 9NQW) | #3 BORF2<br>(EMDB-49688) | #4 A3Bwt<br>(EMDB-49690) |
| --- | --- | --- | --- | --- |
| **Data collection and processing** |  |  |  |  |
| Magnification | 165,000 |  |  |  |
| Voltage (kV) | 300 |  |  |  |
| Electron exposure (e-/Å<sup>2</sup>) | 50 |  |  |  |
| Defocus range (μm) | -0.75 to -2.5 |  |  |  |
| Pixel size (Å) | 0.7296 |  |  |  |
| Symmetry imposed | C2 | C1 | C1 | C1 |
| Initial particle images (no.) | 7,480,511 |  |  |  |
| Final particle images (no.) | 84,950 | 87,889 | 169,180 | 87,889 |
| Map resolution (Å) | 2.67 | 2.61 | 2.42 | 2.81 |
| FSC threshold | 0.143 | 0.143 | 0.143 | 0.143 |
| Map resolution range (Å) |  |  |  |  |
| **Refinement** |  |  |  |  |
| Initial model used (PDB code) | 7rw6 |  |  |  |
| Model resolution (Å) | 3.2 |  |  |  |
| FSC threshold | 0.143 |  |  |  |
| Model resolution range (Å) |  |  |  |  |
| Map sharpening *B* factor (Å<sup>2</sup>) | 71.1 |  |  |  |
| Model composition |  |  |  |  |
| Non-hydrogen atoms | 17608 |  |  |  |
| Protein residues | 2130 |  |  |  |
| Ligands | 10 |  |  |  |
| *B* factors (Å<sup>2</sup>) |  |  |  |  |
| Protein | 57.63 |  |  |  |
| Ligand | 68.58 |  |  |  |
| R.m.s. deviations |  |  |  |  |
| Bond lengths (Å) | 0.004 |  |  |  |
| Bond angles (°) | 1.020 |  |  |  |
| Validation |  |  |  |  |
| MolProbity score | 1.96 |  |  |  |
| Clashscore | 7.99 |  |  |  |
| Poor rotamers (%) | 3.24 |  |  |  |
| Ramachandran plot |  |  |  |  |
| Favored (%) | 0.09 |  |  |  |
| Allowed (%) | 2.65 |  |  |  |
| Disallowed (%) | 97.25 |  |  |  |

The central interaction surface within the A3Bwt-BORF2 complex is the BORF2 dimerization interface (**Figure 2c**). The BORF2 dimer interface is mediated by anti-parallel packing of two α-helices, α12 and α13 (**Figure 2d**). Helix α12 from the juxtaposed BORF2 protomers (chains B and D in the PDB) forms interdigitated aromatic side-chain stacking, involving Trp198, Tyr205, and Trp213 from both chains. In comparison, helices α12 and α13 were mostly disordered in our previously reported A3Bctd-BORF2 complex^18^ (**Figure 2e**). Our prior A3Bctd-BORF2 complex, like the A3Bwt-BORF2 complex reported here, was tetrameric with two copies of each protein. However, this prior complex was nucleated by a non-canonical BORF2 dimerization interface mediated by the N-terminal α-helices of BORF2. This difference could be due to the exclusive use of A3B-CTD in the prior structure, consistent with a role for the A3B-NTD in stabilizing the BORF2 dimer interface observed here. Cellular RNRα subunits from bacteria and yeast to rodents and humans use the same canonical α-helices, corresponding to α12 and α13 of BORF2, for dimerization^34^ (**Figure 2f**). Cellular RNRα subunits also have a regulatory site called the selectivity site, which is formed by a loop region between β2 and α13 (ref.^34^; **Figure 2e,f**). This site binds to dNTPs and helps to establish active site selectivity for specific NDPs. In BORF2, the corresponding loop region is truncated, which likely blocks NTPs from binding and prevents allosteric regulation of the active site.

Interestingly, additional cryo-EM map density was visible on the periphery of the tetrameric A3Bwt-BORF2 complex (**Figure 2g**). This density was explained by an additional BORF2 protein binding to the complex through the non-canonical BORF2 dimerization interface (**Figure 2e**). The presence of both canonical and non-canonical BORF2 dimers in the same cryoEM reconstruction suggests that higher-order oligomers of A3Bwt-BORF2 may be mechanistically viable. BORF2-mediated polymerization of A3B is consistent with work showing that A3Bctd colocalizes with BORF2 in cytoplasmic aggregates in living cells and, in some instances, can appear with a filament-like morphology^15,17,18^. These cytoplasmic structures appear significantly different for full-length A3Bwt-BORF2 complexes^15^, which may be due to formation of a different higher-order oligomer and/or to A3B-NTD binding to RNA and changing the overall nature of the higher-order assembly.

### A3Bwt structure

The A3Bwt-BORF2 complex visualized by cryoEM reveals the first structure of full-length A3Bwt in any context. The pseudo-active NTD and the active CTD of A3Bwt are oriented linearly in a head-to-tail fashion, defined by a 101-degree rotation along an axis connecting the center of each domain (**Figure 3a**). This structure has several notable features (**Figure 3b**). First, the A3Bwt-BORF2 structure shows the CTD interacting with BORF2 through loops 1 and 7, which blocks deaminase activity and confirms our prior studies^18^. However, our new A3Bwt-BORF2 cryoEM map enabled us to model CTD loop 1 differently by positioning Arg211, part of a triple arginine motif in loop 1 (Arg210-Arg211-Arg212), such that its side chain points back towards the core of the CTD rather than outward toward BORF2 (**Figure 3c,d**). This path agrees with prior crystal structures of DNA-free A3Bctd^35,36^ (**Figure 3d**) and suggests BORF2 can accommodate multiple loop 1 conformations. Second, A3B itself forms a homodimeric interface in the complex, mediated by a hydrophobic patch near CTD loop 2 comprised of Leu223, Trp228, and Ile272 (**Figure 3b,d,e**). Gly226 from the opposite A3Bctd protomer butts up against this hydrophobic patch. Third, A3Bntd makes an additional contact with the opposite BORF2 molecule (chain D) from the one bound to A3Bctd (chain B) (**Figure 3b,d,f**). This interaction occurs between loop 4 of A3Bntd (residues 82-85) and residues 546-552 of BORF2. As only ∼166 Å^2^ of surface area is involved in this interaction, it may be weak compared to the A3Bctd-BORF2 interface, which inhibits A3B catalytic activity and buries 1137 Å^2^. In addition, the loop 4 density is weaker compared to other areas of A3Bntd and disappears at higher contour levels, suggesting intrinsic flexibility. Last, the A3Bntd loop 2 (residues 42-45) extends toward an acidic patch on BORF2 formed by residues 377-381 and 322-329, but the density for this loop disappears at higher contour levels, and this contact buries 119 Å^2^ of surface area, also suggesting a relatively weak interaction. Altogether, these interactions are likely to combine to generate an avidity effect that promotes the formation of higher-order assemblies observed here (**Figure 1**) and in cells (refs.^15,17,18^ and see below).

**Figure 3:**
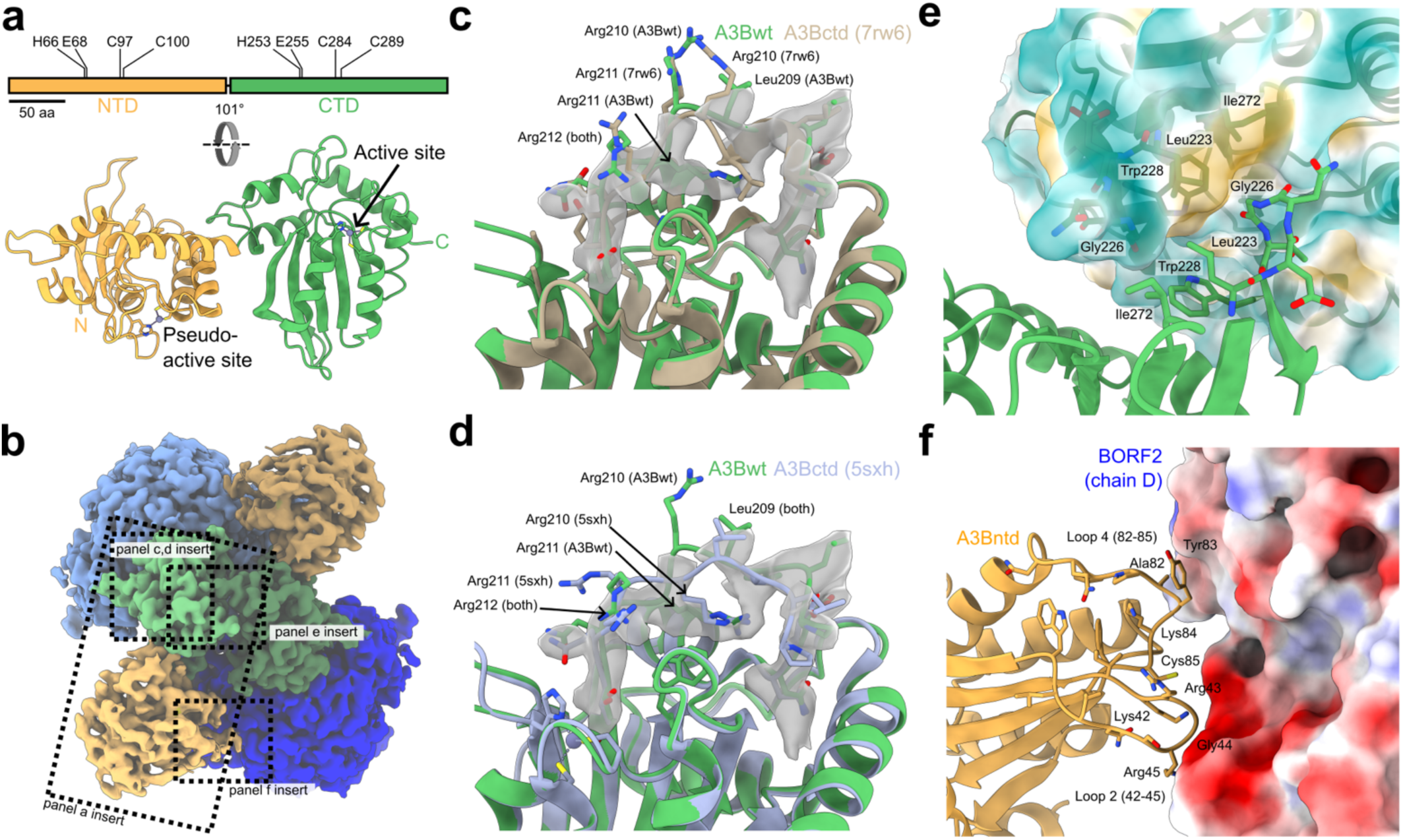
The full-length structure of A3Bwt. **a**, Linear (top) and ribbon (bottom) schematics of A3Bwt with NTD (Z2) and CTD (Z1) domains colored (orange and green, respectively) and zinc-coordinating residues indicated (scale = 50 residues). The domains are rotated 101° relative to each other around an axis connecting the center of both domains. **b**, CryoEM density of the heterotetrameric A3Bwt-BORF2 complex with BORF2 colored as in Figure 2 and A3B colored orange and green as in panel a. Dashed boxes indicate regions expanded in panels a and c-f. **c-d**, Overlay of the active site of A3Bwt (green, this study) with the A3Bctd from the A3Bctd-BORF2 cryoEM structure^18^ (pdb: 7rw6, tan) and an A3Bctd crystal structure^36^ (pdb: 5sxh, lavender), respectively. Loop 1 residues are shown as sticks, and segmented cryoEM density for loop 1 is shown as transparent gray. **e**, Ribbon diagram of the A3Bctd-A3Bctd dimer interface with opposing chains colored as in panel b. A transparent surface representation of chain C is shown colored by hydrophobicity (hydrophobic = orange, hydrophilic = green). Side chains of Leu223, Gly226, Trp228, and Ile272 are shown as sticks. **f**, Structural representation of the A3Bntd-BORF2 interface with A3Bntd (orange) shown as ribbons with loop 2 and 4 side chains as sticks. Chain D of BORF2 is shown in surface representation, colored by electrostatic potential (positive = blue, negative = red).

### A3Bwt hinge region

The A3Bwt-BORF2 cryoEM complex also revealed multiple “hinge region” contacts between the globular NTD and CTD domains (**Figure 4a**). Specifically, the side chains of A3B-NTD Arg133, Gln140, and Tyr191 bridge the NTD-CTD interface by hydrogen-bonding to protein backbone atoms from A3Bctd (**Figure 4a**). This interdomain interaction is further stabilized by CTD Tyr350, which fits into a shallow hydrophobic pocket on the NTD surface and is hydrogen-bonded to Ser139.

**Figure 4:**
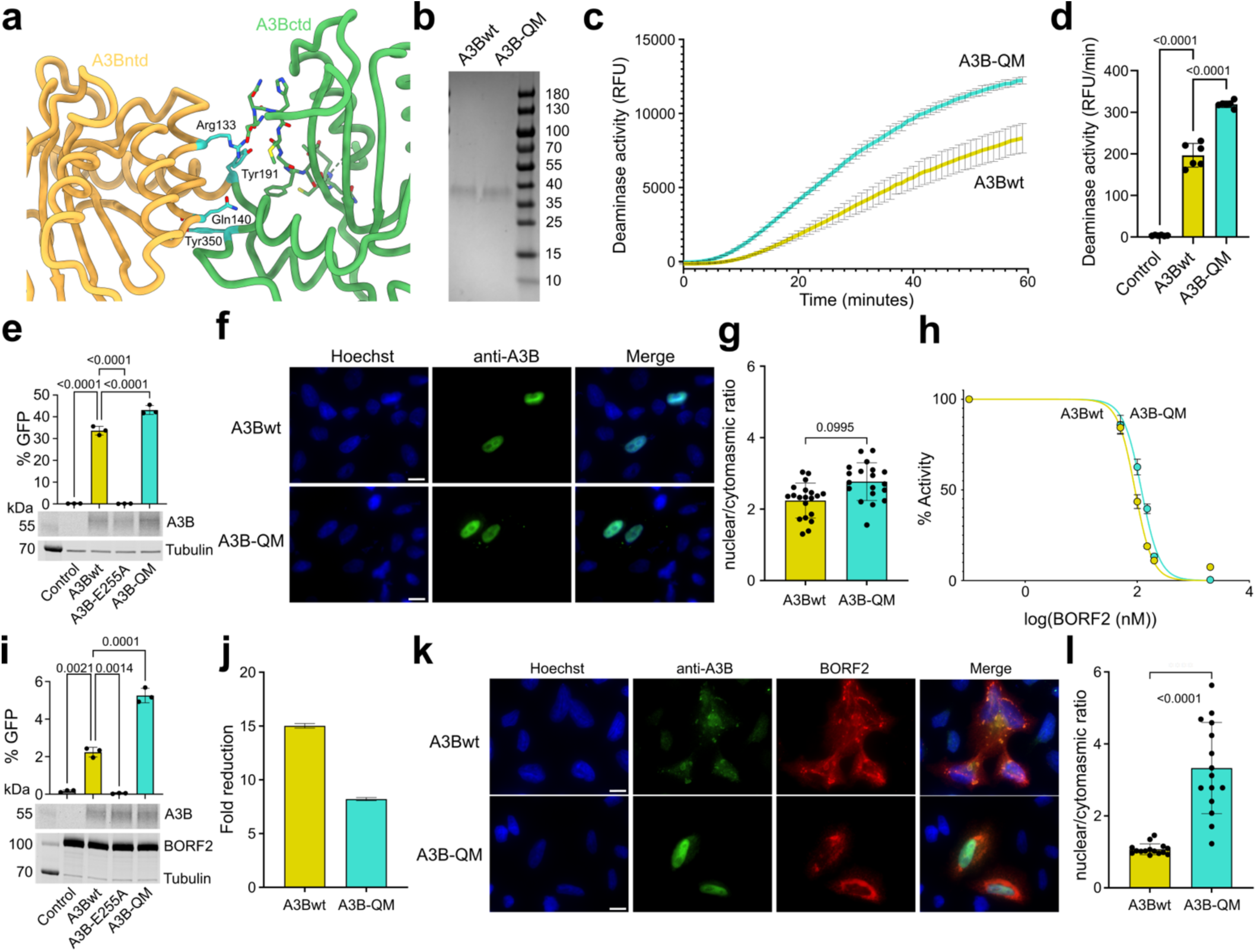
NTD-CTD interaction modulates the A3B activity and sequestration by BORF2. **a**, Structural analysis of the A3Bwt hinge region. Interacting residues Arg133, Gln140, Tyr191, and Tyr350 are shown as sticks and colored cyan. Strand 2 of A3Bctd is also shown in stick format. **b**, Coomassie-stained SDS-PAGE image of A3Bwt and A3B-QM used in DNA deamination experiments. **c-d**, Representative DNA deamination reaction trajectories and rate quantification, respectively, for A3Bwt and A3B-QM in the RADD assay. Data represent the mean ± SD of 6 biological replicates, with some error bars smaller than the data points (*P*-values, one-way Anova test). **e**, DNA deaminase assay in living cells. Top, eGFP fluorescence of 293T cells co-transfected with the AMBER reporter and A3Bwt, A3B-E255A, or A3B-QM plasmid. Data represent mean ± SD of 3 biological replicates, with some error bars smaller than the data points (*P*-values, one-way Anova test). Bottom, immunoblots confirm similar expression levels between the constructs with tubulin as a loading control. **f-g**, Fluorescence microscopy images and quantification of A3Bwt and A3B-QM expressed in HeLa cells. A3B (green) was detected with the 5210-87-13 mAb^45^ (1:300) and nuclei with Hoechst (blue). Data represent the mean ± SD of 15-20 cells/condition (*P*-values, one-way Anova test). Scale bar is 10 µm. **h**, Dose-response plot for BORF2 inhibition of A3Bwt and A3B-QM in the RADD assay. Assays were done with 180 nM of A3B or A3B-QM, and IC_50_ values are 91 nM (95% CI: 86-96 nM) and 122 nM (95% CI: 116-128 nM), respectively. **i**, BORF2 inhibition of A3B DNA editing activity in living cells. Top, eGFP fluorescence of 293T cells co-transfected with the AMBER reporter, BORF2 plasmid, and A3Bwt, A3B-E255A, or A3B-QM plasmid. Data represent the mean ± SD of 3 biological replicates, with some error bars smaller than the data points (*P*-values, one-way Anova test). Bottom, immunoblots confirm similar expression levels between the constructs. **j**, Quantification of the fold-reduction in % eGFP-positive cells upon BORF2 co-expression in the AMBER assay. Data represent the mean ± SD of experiments in panels e and i, which were done at the same time but presented separately for clarity (*P*-value, one-way Anova test). **k**, Fluorescence microscopy images of A3Bwt and A3B-QM co-expressed in HeLa cells with EBV BORF2 (red). A3B (green) was detected with the 5210-87-13 mAb^45^ (1:300), FLAG-tagged BORF2 with a mouse anti-FLAG-M2 antibody (Sigma-Aldrich; 1:500), nuclei with Hoechst (blue). Scale bar is 10 µm. **l**, Quantification of BORF2-mediated relocalization of A3Bwt and A3B-QM in HeLa cells. Nuclei were gated based on the Hoechst staining and the ratio of the mean nuclear to cytoplasmic signal was determined. Data represent the mean ± SD of 15-20 cells/condition (*P*-values, one-way Anova test).

The functional significance of the A3B NTD-CTD hinge region interaction was tested by comparing A3Bwt and a quadruple mutant derivative (A3B-QM = Arg133Ala/Gln140Ala/Tyr191Ala/Tyr350Ala) in multiple assays. First, these two proteins were purified and tested for ssDNA deaminase activity using the RADD assay^32,33^ (**Figure 4b**). Interestingly, A3B-QM exhibited 62% higher rates of ssDNA deamination than A3Bwt (representative kinetics in **Figure 4c** and rate quantification in **Figure 4d**). Second, to test if hyperactivity may be generalizable beyond *in vitro* conditions, the activities of A3Bwt and A3B-QM were compared in living cells using an eGFP DNA editing reporter construct (AMBER), where deamination of a single C-to-U leads to a C-to-T mutation and restoration of eGFP function^37,38^. In comparison to vector control and A3B-E255A catalytic mutant expressing cells, 293T cells transfected with A3Bwt or A3B-QM constructs exhibit high levels of DNA C-to-U editing activity as evidenced by 31.8 and 40.2% eGFP-positive cells after a 48-hour incubation (**Figure 4e**, bar graph above). Third, this highly significant 26% increase in A3B-QM editing activity in the cellular DNA editing assay is consistent with the biochemical results above and unlikely to be due to expression differences, as the two proteins were detected at similar levels (**Figure 4e**, immunoblots below). Fourth, A3Bwt and A3B-QM constructs were examined at the single-cell level by immunofluorescent (IF) microscopy. These experiments showed that both proteins are expressed at similar levels in HeLa cells and, importantly, localize to the nuclear compartment (representative images in **Figure 4f** and quantification in **Figure 4g**), consistent with prior reports^11–14^. Altogether, these results indicate that the loss of the A3B hinge region interaction triggers an intrinsic change in the full-length protein that manifests as elevated DNA deamination activity.

### BORF2-mediated relocalization of A3Bwt and quadruple mutant

Previous work has shown that BORF2 sequesters A3B in cytoplasmic aggregates that can show fibrous morphologies^15,17,18^. As described above, our new cryoEM structure provides a molecular explanation for such higher-order structures, supported by both the canonical and non-canonical dimerization interfaces of BORF2, the BORF2-A3Bctd interface, the BORF2-A3Bntd interface, and the A3B-CTD dimerization interface. The extensive network of interactions suggests that the domain arrangement and specific conformation of A3Bwt may be important for its efficient cytoplasmic sequestration and aggregation by BORF2. In other words, such an assembly can extend in multiple directions to “soak-up” free A3Bwt, provided BORF2 is expressed in molar excess during lytic viral DNA replication.

To investigate this “sponge” model, the A3B inhibition activity of BORF2 was first compared for A3Bwt and A3B-QM. As expected, BORF2 showed a concentration-dependent inhibition of 180 nM A3Bwt in the RADD assay, with an IC_50_ of 91 nM (95% CI: 86-96 nM), demonstrating a tight-binding, stoichiometric inhibition (quantification in **Figure 4h**). In comparison, A3B-QM reproducibly showed a partial resistance to the inhibitory activity of BORF2, with an IC_50_ of 122 nM (95% CI: 116-128 nM). Although this drop in inhibitory potency may seem relatively modest, the IC_50_ value being significantly greater than 50% of the A3B concentration in the assay clearly shows a compromised inhibition of A3B-QM by BORF2. This result suggests that the conformation, and perhaps the rigidity, of A3Bwt may be important for binding and inhibition by BORF2.

To further test this idea, the inhibition, binding, and relocalization of A3B by BORF2 were tested in living cells. A3B-QM was inhibited less by BORF2 in the AMBER assay, with an 8-fold reduction in activity compared to a 15-fold reduction for A3Bwt (**Figure 4i-j**). A3Bwt shows strong nuclear localization, and BORF2 promotes its relocalization out of the nucleus, leading to a reduction in the nuclear/cytoplasmic ratio of A3Bwt (representative images in **Figure 4k** and quantification in **Figure 4l**, mirroring our prior studies^11–14^). By contrast, A3B-QM, which showed similar nuclear localization to A3Bwt in the absence of BORF2 (above), remained enriched in the nucleus in the presence of BORF2 and exhibited little change in the nuclear/cytoplasmic ratio (**Figure 4k-l**). These results further indicate that BORF2 is less capable of binding to and inhibiting A3B-QM, which culminates in a dramatic deficiency in enzyme relocalization.

### A3Bwt structural orientation shows significant differences with A3G

Having validated the A3Bwt structure presented here, we next compared the interdomain organization of A3Bwt (this study), A3Bwt predicted by AlphaFold2 (refs.^39,40^), and A3Gwt from recent work^41,42^. A3B and A3G are closely related enzymes, as the only double-domain deaminases in humans with a Z2-Z1 zinc-coordinating domain organization^6,7^. The predicted A3Bwt structure in the AlphaFold database (A3B-AF), has low confidence (greater PAE: predicted aligned error) for relative positions of residues from A3Bntd and A3Bctd. It is, therefore, not surprising that after aligning A3Bctd, A3Bntd-AF shows an approximately 180-degree rotation from our A3Bntd position, about the axis that connects the centers of the NTD and CTD (**Figure 5a**).

**Figure 5:**
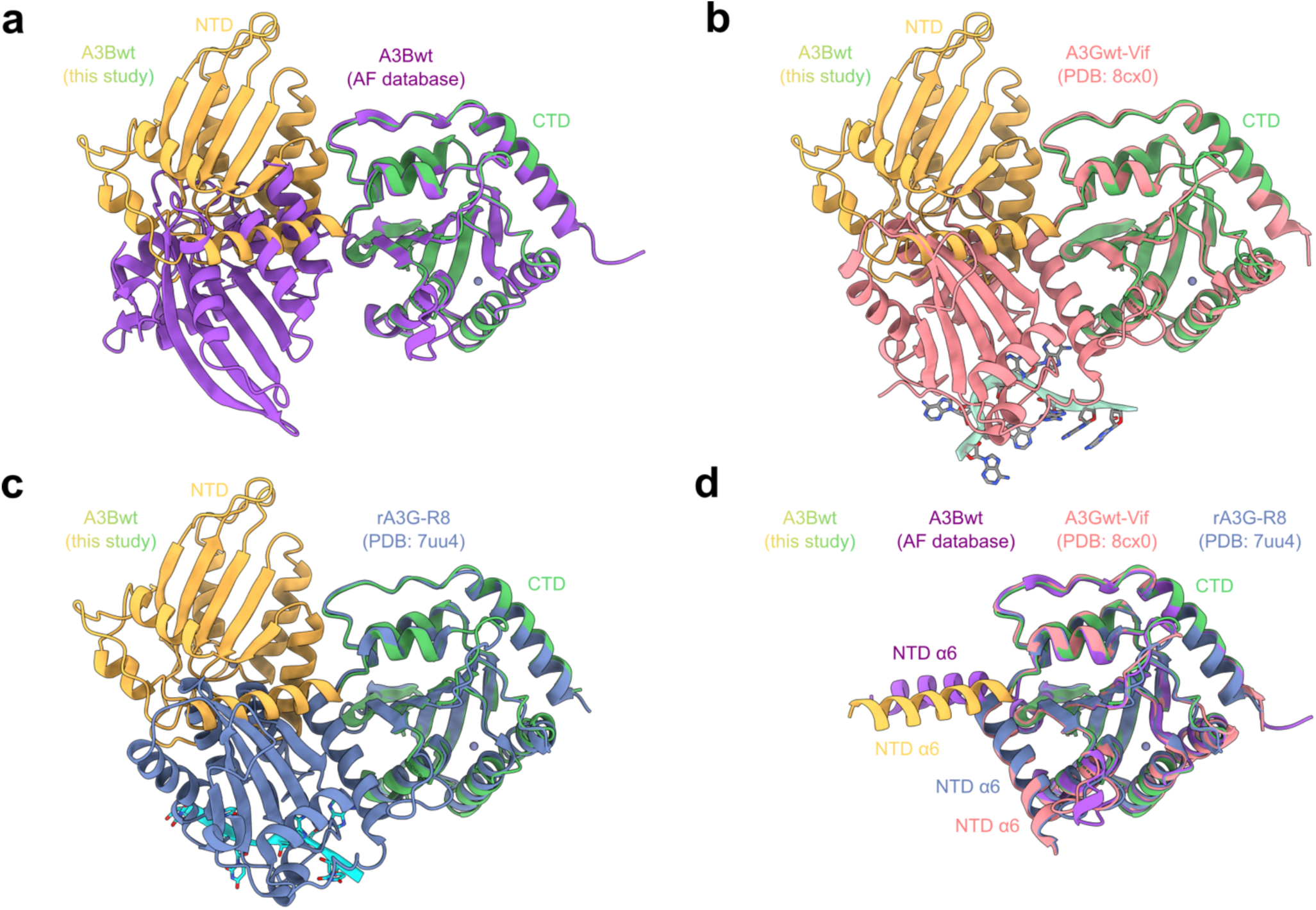
A3Bwt exhibits a unique domain arrangement. Structural analysis of the domain arrangement of A3Bwt (this study) with A3Bwt-AF (**a**, purple), A3Gwt in complex with ssRNA and Vif ^41^ (**b**, pdb: 8cx0, pink), and rA3G-R8 in complex with ssRNA ^43^ (**c**, pdb: 7uu4, blue). A3Bwt is colored as in Figure 3. The CTDs of each structure was aligned to A3Bctd. **d**, Comparison of the structures in **a-c** where only α-helix 6 of NTD is shown to highlight the differences in the domain positioning.

The structure of full-length wildtype human A3G in complex with ssRNA, HIV-1 Vif, CBFβ, ELOB, and ELOC, and that of rhesus macaque A3G (with its NTD loop 8 exchanged to the cognate sequence from human A3B; rA3G_R8_) in complex with ssRNA were reported recently (refs.^41–43^). Beginning with A3Gwt in complex with Vif, after aligning A3Gctd to A3Bctd, a large rotation of ∼90 degrees of A3Gntd to A3Bntd is observed (**Figure 5b**). rA3G_R8_ bound to ssRNA shows an identical conformation to A3Gwt, suggesting that Vif does not induce structural changes in A3G (**Figure 5c**). The magnitude of these differences is highlighted by the position of the last α-helix (helix 6) of the NTD, which projects directly away from the CTD in our A3Bwt structure (**Figure 5d**). In the AlphaFold database structure, helix 6 rotates away from the CTD active site. In the A3G structures, helix 6 of the NTD is cradled by loop 3 and α-helix 2 of the CTD, burying ∼850 Å^2^ of surface area between the helix 6 of the NTD and the CTD compared to only ∼200 Å^2^ in A3B. Even though A3G does not bind to BORF2, the structures of A3G and the AlphaFold predicted structure of A3B were superimposed onto the A3Bwt-BORF2 complex to determine if their domain arrangements could be accommodated without causing steric clashes (**Extended Data Figure 3**). The CTD of these models was aligned to that of A3Bwt due to the higher affinity A3Bctd-BORF2 interaction. No steric clashes were identified in this modeling exercise, providing further evidence that the domain positioning of A3Bwt observed here is not dictated by BORF2 binding, but rather it is likely to be intrinsic to A3Bwt itself. Together with our cryoEM structure of A3Bwt and mutational analyses of the A3Bwt hinge region, these results combine to support the likelihood that the NTD and CTD halves of A3B and A3G have distinct structural orientations and inter-domain contacts, likely reflective of different biological substrates.

## Discussion

Here, the first structure of the wildtype, full-length human A3B is presented. The poor solubility of A3Bwt was overcome and a natural viral antagonist of A3B, EBV BORF2, was leveraged to render A3Bwt amenable to cryoEM analysis. This 278 kDa host-pathogen complex is comprised of two copies of each protein held together by three interfaces: the canonical dimerization interfaces of BORF2 (higher affinity) and separate interactions between A3Bctd and A3Bntd with different protomers of BORF2 (lower affinity). The overall structure of A3Bwt, and particularly the unique interdomain orientation (hinge region), was validated by DNA deaminase activity assays and subcellular localization studies. Most importantly, disruption of the unique NTD-CTD hinge region interaction leads to a hyper-active enzyme, potentially due to an improved motility of the catalytic CTD and therefore an increased capacity to access single-stranded DNA substrate cytosines. Moreover, the unique interdomain orientation of full-length A3Bwt is distinct from that of human A3G (the most closely related human double-domain deaminase), which is likely related to biological function in innate immunity, with A3B strongly linked to restriction of herpesviruses and A3G equally strongly to retroviruses.

To our surprise, A3B adopts a completely different interdomain arrangement than A3G, even though the proteins have similar size (382 and 384 aa), conserved zinc-coordinating domains (Z2 and Z1), and high amino acid identity (50% NTD and 64% CTD). These differences are likely to reflect known mechanistic differences between the proteins. Binding of A3G to RNA regulates cytoplasmic localization, self-oligomerization, incorporation into HIV-1 virus particles, and binding by HIV-1 Vif. A3B is nuclear localized and capable of organization into high molecular weight aggregates, which reduces its deaminase activity. A low intrinsic activity may be important to limit mutation of the host genome when no innate immune response is occurring, though presumably it must somehow be activated during lytic herpesvirus viral infections.

The A3Bwt-BORF2 heterotetrameric complex characterized here by cryoEM also suggests a mechanism for A3B neutralization by EBV BORF2. The canonical and non-canonical BORF2 dimerization interfaces provide interaction surfaces for nucleating a much larger superstructure (**Extended Data Figure 4**). These interactions alone can nucleate the formation of sponge-like assemblies comprised of canonical dimers linked together through the non-canonical dimerization surface. The A3Bwt-BORF2 heterotetrameric complex shown here strongly indicates that the canonical BORF2 dimer is likely to be stabilized by binding to two A3Bwt protomers, with the NTD of one protomer binding directly to one copy of BORF2 and the CTD more strongly to the other. This tetrameric complex is then further stabilized by CTD-CTD interactions of two bound A3Bwt enzymes. Together, the formation of the larger BORF2 superstructure provides an elegant mechanism to literally “soak-up” cellular A3Bwt and effectively neutralize its antiviral activity. Such a model is consistent with the large cytoplasmic aggregates observed here (Figure 4k) and previously^15–17^ with A3Bwt-BORF2 in cells.

As a major source of mutagenesis in cancer^19–25^, A3B is a target for therapeutic inhibition. Like the profound inhibition of A3B by BORF2 and related herpesviral RNRα subunits^15,17,18,44^, small molecule inhibitors (or degraders) of A3B would be predicted to protect chromosomal DNA from A3B-catalyzed deamination and mutagenesis. Such inhibitors of tumor evolvability would make tumor genomes more stable and less prone to developing drug-resistance mutations and becoming metastatic. The full-length A3Bwt structure provided by our studies (enabled by the BORF2 structural platform) could facilitate A3B inhibitor development directly (BORF2 mimics) and indirectly by enabling high-resolution modeling of candidate inhibitors in the context of the wildtype enzyme.

## Methods

### Solubility testing

Expi293F™ cells (Thermo Fisher Scientific) were seeded at a density of 3 million viable cells/mL in 500 ml of fresh Expi293™ expression media. 500 µg pcDNA3.1-A3Bi-mycHis^23,32^ was diluted in Opti-MEM™ I Medium with 1.6 ml of polyethylenimine (PEI, 10 mg/ml) and incubated for 20 mins. The complex was slowly added to the cells, then incubated at 37^°^C and 8% CO_2_ on an orbital shaker. After 18–22 hrs, valproic acid (4.4 mM), sodium propionate (6.1 mM), glucose (40 mM), non-essential amino acids, and an anti-clumping agent (Thermo Fisher Scientific) were added to the cells to enhance protein expression. The cells were harvested 3 days after transfection and washed with PBS. A 2X base buffer was formulated containing 50 mM Tris-HCl, pH 8.0, 1 M NaCl, 10 mM MgCl_2_, 40 mM imidazole, 10% glycerol, 10mM β-mercaptoethanol (BME). Two cOmplete EDTA-free protease inhibitor cocktail tablets (Roche) were added to 50 mL of buffer for a final 2X concentration. For detergent additives, IGEPAL CA-630, 3-((3-cholamidopropyl) dimethylammonio)-1-propanesulfonate (CHAPS), TERGITOL (NP-40), and Triton X-100, 10% solutions were created. For other additives, 10X solutions were created (500 mM L-arginine, 500 mM L-proline, 500 mM imidazole), apart from L-glutamine, which was formulated to 250 mM due to solubility limitations. The pH of the buffer was adjusted using 6 M NaOH if necessary. Cell aliquots were first thawed before adding 500 μL 2X base buffer. 10 μL 10% detergent mixtures were added to respective samples for a final concentration of 0.1%, followed by 100 μL 10X additives (200 μL for L-glutamine). All sample volumes were then brought to a final volume of 1 mL with sterile water. Samples were then rocked at 4 ^°^C for 1.5 hrs. Lysate samples were briefly mixed before adding 0.5 mg (2.5 μL) PureLink RNase A (Thermo Fisher Scientific) and 12.5 units (0.5 μL) Salt-Active Nuclease (SAN) (Sigma-Aldrich). Lysate samples were then incubated at 37 ^°^C for 45 mins. An aliquot of total lysate was taken from each sample and stored at 4 ^°^C before the remaining samples were centrifuged at 16,000 xg at 4^°^C for 30 mins. Clarified lysate samples were then removed with care taken to ensure the insoluble pellet was not disturbed. Total lysate and clarified lysate samples were then diluted 1:10 in SDS-PAGE loading buffer before electrophoresis. The resulting gel was then transferred to a membrane, which was blocked overnight at 4^°^C in 4% milk + PBST. Primary blotting was performed with 1:5,000 rabbit anti-A3B 5210-87-13 (ref.^45^; Cell Signaling) and 1:5,000 mouse anti-alpha-tubulin T5168-100UL (Sigma-Aldrich) in PBST for 2 hrs at room temperature (RT). Secondary blotting was performed using 1:20,000 680LT goat anti-rabbit 925-68021 (LI-COR) and 1:20,000 800CW goat anti-mouse 926-32210 (LI-COR) in PBST + 0.01% SDS for 2 hrs at RT. Membranes were imaged using an Amersham Typhoon biomolecular imager using the built-in IRshort and IRlong methods.

### Full-length APOBEC3B expression and purification

Full-scale purification followed the same procedure as above. Cells were harvested 72 hrs post-transfection, pelleted by centrifugation for 5 mins at 1000 x g, split evenly into two pellets, and then frozen and stored at −80 ^°^C. The two pellets were resuspended in 40 mL lysis buffer consisting of 25 mM Tris-HCl pH 9.0, 500 mM NaCl, 5 mM MgCl_2_, 20 mM imidazole, 5% Glycerol, 0.1% IGEPAL CA-630, 50 mM L-arginine, 5mM BME, and one cOmplete EDTA-free protease inhibitor cocktail tablet (Roche) per 50 mL buffer. The lysis buffer was adjusted to a final pH of 9.0 with NaOH and filter sterilized with a 0.22 μm membrane. Cell pellets were disrupted via pipetting until no visible clumps remained. The lysis mixture was then sonicated for 2 mins on ice with a Branson Analog Sonifier model 450 (50% duty cycle – output level 5) and incubated on ice for 2 mins before repeating the sonication cycle. 5 mg (250 µL) PureLink RNase A (Thermo Fisher Scientific) and 5000 U (50 µL) Salt-Active Nuclease (NEB) were added to each tube (∼50 mL lysate per tube), and the lysate was then incubated at 37 ^°^C for 1.5 hrs. After incubation, the lysate was brought up to a final NaCl concentration of 1 M and centrifuged at 20,000 x g at 4 ^°^C for 30 mins. Clarified lysate was vacuum filtered using a 0.22 μm membrane. A 5 mL Ni-NTA Superflow column (Qiagen) cartridge was pre-equilibrated with lysis buffer before loading the clarified lysate at a flow rate of 0.5 mL/min. The column was then washed at 0.5 mL/min with high-salt wash buffer containing 25 mM Tris-HCl, pH 9.0, 1 M NaCl, 20 mM Imidazole, 10% Glycerol, 0.1% CHAPS, and 5 mM BME until the UV signal returned to baseline. The protein was then washed with 50 mL of reduced-salt buffer containing 25 mM Tris-HCl, pH 9.0, 500 mM NaCl, 20 mM Imidazole, 10% Glycerol, 0.1% CHAPS, and 5 mM BME at a flow rate of 0.5 mL/min. The protein was eluted with 100% elution buffer containing 25mM Tris-HCl, pH 9.0, 500 mM NaCl, 400 mM Imidazole, 10% Glycerol, 0.1% CHAPS, and 5 mM BME in 2 mL fractions over a total volume of 100 mL at a flow rate of 0.5 mL/min. Protein was generally eluted from the column between 5 and 35 mL after 100% elution buffer flow began; however, A3B was detectable until ∼90 mL. Fraction content was verified via SDS-PAGE before being concentrated to a volume of ∼5 mL using an Amicon Ultra-15, 10,000 Da MWCO centrifugal concentrator. Following concentration, the protein immediately underwent size-exclusion chromatography by injection into an NGC Quest 10 Plus system equipped with a Hiload™ 26/600 Superdex 75 pg column (320 mL bed volume). The sample was separated using a buffer comprised of 25 mM Tris-HCl pH 9.0, 500 mM NaCl, 10% Glycerol, 0.1% CHAPS, and 0.5 mM Tris (2-carboxyethyl) phosphine (TCEP) at a flow rate of 0.2 mL/min to achieve maximum separation. Fraction contents were confirmed via SDS-PAGE and separated into two pools based on purity, concentrated to ∼1 mL using Amicon Ultra-15, 10,000 Da MWCO centrifugal concentrators, aliquoted, and snap-frozen in liquid nitrogen. All samples were stored at −80 ^°^C.

### Epstein-Barr Virus BORF2 (1-738) expression and purification

The BORF2 construct tagged with a maltose-binding protein (MBP) and a 6x His-tag [pmalx(E)-BORF2(1-738)] was transformed into competent BL21(DE3) *E. coli* before plating onto an LB-agar plate containing 100 µg / mL carbenicillin. Colonies were grown overnight at 37 ^°^C before being resuspended in sterile water and distributed into 4 1-liter flasks of Miller’s LB broth containing 100 μg/ml Ampicillin. The cultures were grown at 37 ^°^C with shaking at 200 rpm until an OD600 of ∼1.0 was reached. The cultures were then induced with 0.5 mM Isopropyl β-D-1-thiogalactopyranoside (IPTG) and incubated at 18 ^°^C for 18 hrs. The cells were pelleted via centrifugation at 4,000 x g for 30 mins and resuspended in lysis buffer containing 20 mM Tris-HCl, pH 7.4, 500 mM NaCl, 5 mM imidazole, and 5 mM BME. Cells were lysed with the addition of 40 mg hen egg white lysozyme followed by sonication (6 cycles, 30 pulses/cycle, 50% duty cycle – output level 5). The cell lysate was centrifuged at 64,000 x g, at 4 ^°^C for 1 hr, followed by vacuum filtration through a 0.22 μm membrane. The filtered lysate was loaded into a 5 mL Ni-NTA Superflow cartridge (Qiagen) equilibrated with lysis buffer at a flow rate of 1 mL/min. Protein elution was carried out with a linear gradient of imidazole by gradually replacing the lysis buffer with an elution buffer containing 50 mM Tris-HCl, pH 7.4, 500 mM NaCl, 400 mM imidazole, 5% glycerol, and 5 mM BME at a flow rate of 2 mL/min, collected in 5 mL fractions. The fractions containing MBP-BORF2(1-738) were confirmed via SDS-PAGE gel and pooled before loading onto a 5 mL MBP-trap cartridge at a rate of 1 mL/min, equilibrated with wash buffer containing 50 mM Tris-HCl pH 8.0, 500 mM NaCl, 10% glycerol, and 1 mM TCEP. The column was washed with buffer until UV signal returned to baseline. MBP-Borf2(1-738) was eluted over 10 × 5 mL fractions at 2 mL/min using 100% elution buffer containing 50 mM Tris-HCl pH 8.0, 500mM NaCl, 10% glycerol, 1 mM TCEP, and 10 mM maltose. The resulting eluate was ∼95% pure and was dialyzed at 4^°^C overnight into a storage buffer containing 20 mM Tris-HCl, pH 8.0, 200 mM NaCl, 1 mM TCEP using a 2,000 Da MWCO Slide-A-Lyzer dialysis cassette (Thermo Fisher Scientific). The sample was then concentrated to a volume of 500 μL using an Amicon Ultra-15, 50,000 Da MWCO spin-concentrator. Protein was flash-frozen in liquid nitrogen and stored at −80 ^°^C.

### Pyrococcus furiosus Endonuclease Q (EndoQ)

EndoQ was purified as previously described with the following modification^33,46^. Due to the thermostability of EndoQ and the need to eliminate a trace amount of co-purified nucleases, the EndoQ stocks were prepared by first heating the enzyme to 60 ^°^C for 15 min and then cooling to RT. Protein concentration was then quantified via NanoDrop and diluted to a final concentration of 20 µM before snap-freezing.

### Protein complex formation and purification

A Superdex 200 increase 10/300 GL column was first equilibrated with buffer containing 20 mM Tris-HCl, pH 8.0, 200 mM NaCl, and 0.5 mM TCEP. A 500 μL injection sample containing 10 μM full-length A3B and 8 μM MBP-BORF2(1-738) was prepared and incubated on ice for 1 hr. Following incubation, the sample was manually injected into a 500 μL sample loading loop and run over the column at a rate of 0.4 mL/min and collected into 0.3 mL fractions. Fraction contents were assayed via SDS-PAGE gel. Fractions containing the full-length A3B – BORF2(1-738) complex were found to elute between 8 and 12 mL. The fractions containing the complex were then concentrated using an Amicon Ultra-0.5 - 3,000 Da MWCO centrifugal filter cartridge to a final volume of 100 μL, resulting in an approximate concentration of 0.5 mg/mL. Samples were immediately used for grid-making and were not frozen or stored.

### CryoEM data collection

3 µL aliquots of 0.5 mg/mL complex were added to UltrAuFoil R1.2/1.3 300 mesh grids (Quantifoil Micro Tools GmbH) loaded into the humidity chamber of a Vitrobot Mark IV (Thermo Fisher Scientific) maintained at 4 ^°^C and 100% humidity. After a ten-second wait, the sample was blotted with a force of −10 for 2-4 seconds and plunged into liquid ethane to vitrify. Grids were shipped to the Pacific Northwest CryoEM Center (PNCC), screened on an Artica TEM, and the best grid was selected for data collection on “Krios 1” at PNCC. 29,146 movies in the EER format were collected with EPU using Image/Beam shift on a Falcon 4i detector at a pixel size of 0.7296 Å/pixel and a total dose of 50 e^-^/Å^2^.

### CryoEM data processing

Movies were imported into CryoSPARC v4.6 and upsampled by a factor of two to a pixel size of 0.3648 Å/pix^47^. Movies were motion corrected using the *Patch Motion Correction* job with an “output F-crop factor” of ¾ (resulting pixel size 0.4864 Å/pix) and with the results saved in 16-bit floating point^48^. The contrast transfer function (CTF) of each aligned movie was determined using the *Patch CTF* job with default settings. All movies collected in the same holes relative to the stage position were grouped using the *Exposure Group Utilities* job to enable beam tilt estimation. Micrographs were denoised using the *Micrograph Denoiser* job with default parameters using a subset of 100 micrographs for training. Micrograph curation was performed with the *Curate Exposures* job resulted in accepting 17.410 micrographs by gating the Average Intensity (0-78.9), CTF fit resolution (Å) (0.974-8), Relative Ice Thickness (1.0-1.2), Total full-frame motion distance (pixels) (6.96-50), and Average defocus (Å) (2000-39058.844), Particles were picked from the denoised micrographs using the *Blob Picker* job with a minimum particle diameter of 100 Å, a maximum particle diameter of 200 Å, and minimum separation distance of 0.5 diameters. The original 7,480,511 particles were curated using a combination of the *Inspect Picks, 2D Classification,* and *Ab-Initio* followed by *Heterogeneous Refinement* jobs. 180,501 particles from accepted 2D classes were input into an *Ab-initio refinement* job with three classes and the best class was selected, duplicate particles were removed and the resulting 91,659 particles were re-extracted with a box size of 600 pixels, Fourier-cropped to 400 pixels (0.7446 Å/pix) and subjected to a *Non-uniform Refinement* job with per-particle defocus estimated and C2 symmetry applied. Particles were polished using a *Reference-based Motion Correction* job and 84,950 particles were re-extracted to the same box and pixel size as the previous refinement job^49^. *Non-uniform Refinement* with no windowing, C2 symmetry, and optimization of both per-particle defocus and per-group CTF parameters was performed^50^. The resulting map at 2.67 Å resolution is the full, tetrameric complex shown in Figure 2b (A3Bwt-BORF2 dimer map, EMDB-49685). Particles were symmetry-expanded and local refined to 2.46 Å with C1 symmetry applied. Focused refinements with masks around BORF2 and A3Bwt yielded maps at 2.42 Å (BORF2 map, EMDB-49688) and 2.77 Å, respectively. The A3Bwt focused refinement went through an additional round of 3D classification with two classes and a filter resolution of 4 Å to select for particles with better A3Bwt occupancy. The resulting 87,889 particles were local refined to 2.81 Å with a mask around A3Bwt (A3Bwt map, EMDB-49690). The same particles were locally refined with a mask around A3Bwt and BORF2 to 2.61 Å (A3Bwt-BORF2 monomer map, EMDB-49687). The data processing pipeline for the A3Bwt-BORF2 dataset is shown in **Extended Data Figure 2**.

### Atomic model building

The A3Bctd-BORF2 complex was used as a starting model for model building^18^. BORF2 was rebuilt in coot using the focus refinement map of BORF2 (ref.^51^). A3Bwt was built in Coot ^52^ using the focus refinement map of A3B. The hA3Bntd structure^30^ was used as a starting point for the NTD with all mutations in that model reverted to wild type. Protein models were real space refined in Phenix^53^. Waters and ions were added to the map using the SWIM plugin in Chimera^54–56^ with Z = 5, threshold of 0.1, water distance 2.5 < 3.1 Å and ion distance 1.9 > 2.5 Å. All ions were manually checked, and those had more nearby atoms at the ideal water hydrogen bonding distance were changed to waters. The resulting model was real space refined again in Phenix. The two models were rigid-body docked into the consensus C2 refined map using Chimera X and adjustments were made in Coot to alleviate any clashes at interfaces between the models^57–59^. A summary of cryoEM data collection, model refinement, and validation statistics is shown in **Table 1**.

### Structural analysis

All structural analysis was carried out in Chimera X^57^. Buried surface area calculations were done using the “measure buriedarea” command. The Alphafold database structure of A3B was downloaded from [https://alphafold.ebi.ac.uk/entry/AF-Q9UH17-F1].

### A3B mutant quantification and concentration normalization

Full-length A3B concentrations were determined based on band intensity in SDS-PAGE, against a dilution series of EndoQ at known concentrations as a reference due to its similar molecular weight. The dilution series was run on an SDS-PAGE gel alongside full-length A3Bwt and A3B-QM. Purified full-length A3B samples were not boiled before being loaded onto SDS-PAGE gels, as doing so causes significant aggregation and results in much of the sample being unable to enter the gel. Gels were stained with SYPRO-Ruby (Invitrogen) using the standard protocol. Gels were scanned with an Amersham Typhoon fluorescence imager using the built-in Cy3 method. The bands were digitally quantified using the Fiji distribution of ImageJ2 and compared to a standard curve for the dilution series of EndoQ to determine the concentration of A3B samples. To verify these results, A3Bwt and A3B-QM were diluted to a uniform concentration, followed by SDS-PAGE and SYPRO-Ruby staining to ensure uniform size and intensity of all bands (**Figure 4b**). These concentration values were used for all deamination assays.

### Single-stranded DNA deamination reporters

The deamination substrate was a 15 nt ssDNA oligonucleotide with a 5′ fluorescein (FAM) (5′-FAM-TAGGT<u>C</u>ATTATTGTG-3’). The single deoxycytidine in the 5′-TC motif is deaminated by A3B to produce a deoxyuridine, which is subsequently cleaved by EndoQ. The real-time DNA deamination assay utilized a modified reporter bearing the same sequence with the addition of a 3′ Iowa Black® FQ (IAB) quencher and an internal ZEN™ quencher (5′-FAM-TAGGT<u>C</u>ATT-ZEN-ATTGTG-IAB-3’). Upon cleavage by EndoQ, the fluorophore is released from the quenchers and produces a signal^32^. Additional oligonucleotides contained a dU in place of the target dC as a control for EndoQ cleavage. All oligonucleotides were synthesized by Integrated DNA Technologies, Inc.

### A3B deamination activity assays and BORF2 inhibition assays

Gel-based deamination assays were prepared as 40 µL reactions containing unquenched reporter (5′-FAM-TAGGT<u>C</u>ATTATTGTG) at a final concentration of 100 nM. For each 40 µL reaction, a reporter master mix contained 4 µL 1 µM Reporter, 4 µL 10X Deamination buffer (600 mM Tris-HCl pH 8.0, 0.1% Tween20, 10% DMSO, 10 mM DTT), 4 µL 50 mM EDTA, and 16 µL Ultra-pure RNase-free water for a total of 28 µL was created. The reporter master mixes were incubated at 60 ^°^C for 15 mins before decreasing the temperature to 37 ^°^C where it stayed until adding enzymes. An enzyme master mix was then prepared in 10X concentration containing 1:1:1 volumes of 1.8 µM A3B, 20 µM EndoQ and BORF2 at varying concentrations. The enzyme master mixes were then incubated at 37 ^°^C for 10 mins to equilibrate them to reaction temperature. After pre-incubation, the enzyme master mixes (12 µL) and reporter master mixes (28 µL) were combined and the reactions were incubated at 37 ^°^C for 1 hr. The reactions were then immediately quenched by the addition of an equal volume of 100% formamide and boiled at 95 ^°^C for 10 mins. Following boiling, an additional reaction volume of 2X xylene cyanol formamide loading buffer was added. 12 µL of sample was then transferred to a 15% TBE-Urea gel and run at 300 V for 25 mins. All gels were imaged with an Amersham Typhoon scanner using the built-in Cy2 method.

Real-time DNA deamination (RADD) assays^32,33,46^ to evaluate the activity of A3Bctd, A3Bwt, and A3B-QM with and without BORF2 were prepared similarly to the 40 µL reactions described above, except the reporter. All RADD reactions utilize the double-quenched fluorescent reporter (5′-FAM-TAGGT<u>C</u>ATT-ZEN-ATTGTG-IAB) at a final concentration of 1 µM. The 10X enzyme master mixes were prepared with 1:1:1 volumes of 1.8 µM A3B, 20 µM EndoQ and RNase-free water. After enzyme and reporter master mix pre-incubation, 12 µL of enzyme master mix for each condition was combined with the reporter master mix and immediately transferred to a pre-warmed 96-well half-well plate and scanned for 1 hr at 37 ^°^C using a TECAN Spark 10M. The final concentrations in this mixture were 1 µM reporter, 180 nM A3B, and 2 µM EndoQ. Each condition was evaluated in 3X technical replicates. For assays evaluating BORF2 inhibition, the reporter master mix was produced identically to the process described above, except the enzyme master mix, which was assembled in 10X concentration containing 1:1:1 ratio of 1.8 µM A3B, 20 µM EndoQ, and BORF2 (variable concentration) or BORF2 storage buffer.

To quantify the activity of full-length wild-type A3B and mutant constructs, the data were first processed to remove background signal. Each active sample was paired with an inactive sample containing reporter and A3B, but lacking EndoQ. The RFU values of these reactions were subtracted from the value of the respective reactions containing reporter, A3B, and EndoQ. Activity was measured by determining the maximum slope achieved by the reaction using the Interactive Continuous Enzyme Analysis Tool (ICEKAT)^60^. The slopes of all replicates were calculated independently. A similar process was utilized to calculate the IC_50_ values for BORF2 inhibition, utilizing ICEKAT to determine the slope of each reaction at a given BORF2 concentration. These slopes were then used to calculate an IC_50_ value by first normalizing the data in Graphpad Prism 9 with the slope of the uninhibited sample representing 100% activity and a slope of 0 RFU/s being 0% activity. The built-in log(inhibitor) vs. normalized method response was then used to calculate the IC_50_ values for all samples.

### A3B activity quantification in cellulo using AMBER

293T cells were seeded in 12-well plates (Corning) and transfected at approximately 85% confluency. Each well received 200 ng AMBER reporter plasmid, 150 ng SpCas9 nickase plasmid (BE4max backbone), 100 ng AMBER gRNA plasmid, 150 ng plasmid encoding the indicated A3B mutant, and 200 ng empty vector or BORF2 expression plasmid. Transfections were performed using 2.4 μL TransIT-LT1 reagent (Mirus Bio, MIR 2300) according to the manufacturer’s instructions. 24 hrs post-transfection, cells were dissociated with trypsin and resuspended in 1 mL complete growth medium. A 200 μL aliquot was analyzed by flow cytometry to quantify mCherry and GFP fluorescence. The remaining cells were collected for immunoblot analysis.

### Immunoblotting of AMBER assays

For protein expression analysis, cells were harvested following collection of aliquots for flow cytometry, washed with phosphate-buffered saline (PBS), lysed in RIPA buffer (Thermo Fisher Scientific, PI89900), and quantified using BCA protein assay (Thermo Fisher Scientific, 23225). Equal amounts of protein were separated by SDS-PAGE on 4-20% Tris-glycine gels (Bio-Rad, 5671095) and transferred to PVDF-FL membranes (MilliporeSigma, IPFL00005). Membranes were washed with PBS containing 0.1% Tween-20 (PBST), blocked in 1× casein blocking buffer (Sigma-Aldrich, C7594), and incubated overnight at 4°C with primary antibodies diluted in blocking buffer. Following three washes with PBST, membranes were incubated for 1 hr at RT with IRDye-conjugated secondary antibodies diluted in blocking buffer supplemented with 0.01% SDS. Membranes were subsequently washed three times with PBST and imaged using a LI-COR Odyssey imaging system. Primary antibodies included rabbit anti-human A3B (5210-87-13; 1:1,000; ref.^45^; Cell Signaling), anti-FLAG (Sigma-Aldrich, F1804; 1:1,000), and anti-α-tubulin (Sigma-Aldrich, T5168; 1:5,000). Secondary antibodies included IRDye 800CW goat anti-rabbit IgG (LI-COR, 827-08365; 1:10,000) and IRDye 680LT goat anti-mouse IgG (LI-COR, 925-68020; 1:10,000).

### Immunofluorescence microscopy and image analysis

HeLa cells were cultured at a density of 50,000 cells per well in 4-chamber, tissue culture-treated glass slides (Falcon). The following day, cells were transiently transfected with 200 ng of A3Bwt or the A3B-QM mutant construct either in the presence or absence of 200 ng of BORF2-FLAG. 48 hrs post-transfection, cells were fixed with 4% paraformaldehyde (PFA) in PBS for 10 mins at room temperature, followed by three washes with PBS. Permeabilization was performed using 0.5% Triton X-100 (Sigma-Aldrich) in PBS for 10 mins at RT, after which the cells were washed three times with PBS. Cells were then incubated in an immunofluorescence blocking buffer (5% goat serum, 2% BSA, and 0.2% Triton X-100 in PBS) for 1 hr at RT. Primary antibody incubation was performed overnight at 4°C using a rabbit anti-human A3A/B/G antibody^45^ (5210-87-13; 1:300) and a mouse anti-FLAG-M2 antibody (Sigma-Aldrich; 1:500) diluted in blocking buffer. The following day, cells were washed 5 times with PBS and incubated with fluorophore-conjugated secondary antibodies—Alexa Fluor 488 goat anti-rabbit (Invitrogen; 1:1,000) and Alexa Fluor 594 goat anti-mouse (Invitrogen; 1:1,000)—in blocking buffer for 1 hr at RT in the dark. Nuclei were counterstained using Hoechst 33342 (Mirus; 1:20,000 in PBS) for 15 min at RT. Following three final washes with PBS, slides were mounted using ProLong™ Glass Antifade Mountant (Invitrogen), allowed to set overnight at RT protected from light, and stored at 4°C. Images were captured at 60× magnification using a Nikon ECLIPSE Ti2 microscope under oil immersion. To evaluate the cellular distribution and abundance of A3B, the nuclear and cytoplasmic intensities of the A3B signal were quantified using Fiji software. Nuclei were defined based on the Hoechst stain boundaries, and the respective nuclear and cytoplasmic regions of interest (ROIs) were segmented to measure the mean fluorescence intensity of the A3B signal per cell. Statistical analyses were conducted in GraphPad Prism 9.

## Acknowledgements

We thank Clare Morris, Mac Kevin Braza, and Rommie Amaro for thoughtful discussions, Nicholas Moeller for assistance in protein purification, and Febri Sugiokto for comments on the manuscript. This work was supported by NIH grants P01-CA234228 to R.S.H. and H.A., R35-GM118047 to H.A., and F31-CA295111 to C.A.B. R.H.A. was supported in part by a mentored project through the Midwest AViDD Center (NIAID 1U19AI171954-01). C.A.B. was supported in part by the UMN Institute of Molecular Virology T32 training program T32AI083196. C.D.M. received salary support from the South Texas Medical Scientist Training Program (NIGMS T32-GM113896 and T32-GM145432) and the Epigenetics, DNA Repair, and Genomics (EDGe) Training Program (NCI T32-CA279363). Cancer research in the Harris lab is also supported by NCI P50-CA247749 and a Recruitment of Established Investigators Award from the Cancer Prevention and Research Institute of Texas (CPRIT RR220053). R.S.H. is an Investigator of the Howard Hughes Medical Institute, a CPRIT Scholar, and the Ewing Halsell President’s Council Distinguished Chair at University of Texas Health San Antonio. A portion of this research was supported by NIH grant R24GM154185 and performed at the Pacific Northwest Center for Cryo-EM (PNCC) on proposal 160269 with assistance from Vamseedhar Rayaprolu. Parts of this work were carried out in the Characterization Facility, University of Minnesota, which receives partial support from the NSF through the MRSEC (Award Number DMR-2011401) and the NNCI (Award Number ECCS-2025124) programs. We thank the Structural Biology Facility in MBiC, The Hormel Foundation, and The Hormel Institute, University of Minnesota, for providing access to a Vitrobot. Molecular graphics and analyses performed with UCSF ChimeraX, developed by the Resource for Biocomputing, Visualization, and Informatics at the University of California, San Francisco, with support from National Institutes of Health R01-GM129325 and the Office of Cyber Infrastructure and Computational Biology, National Institute of Allergy and Infectious Diseases.

## Data Availability

Atomic coordinates for the A3Bwt-BORF2 complex have been deposited in the RCSB Protein Data Bank with the accession code 9NQV (heterotetrameric complex) and 9NQW (heterodimeric complex). CryoEM maps have been deposited in the Electron Microscopy Data Bank (EMDB) with accession codes 49685 (A3Bwt-BORF2 heterotetramer), 49687 (A3Bwt-BORF2 heterodimer), 49688 (BORF2), and 49690 (A3B). All other data and constructs are available from the authors upon request.

---

**Extended Data Figure 1:**
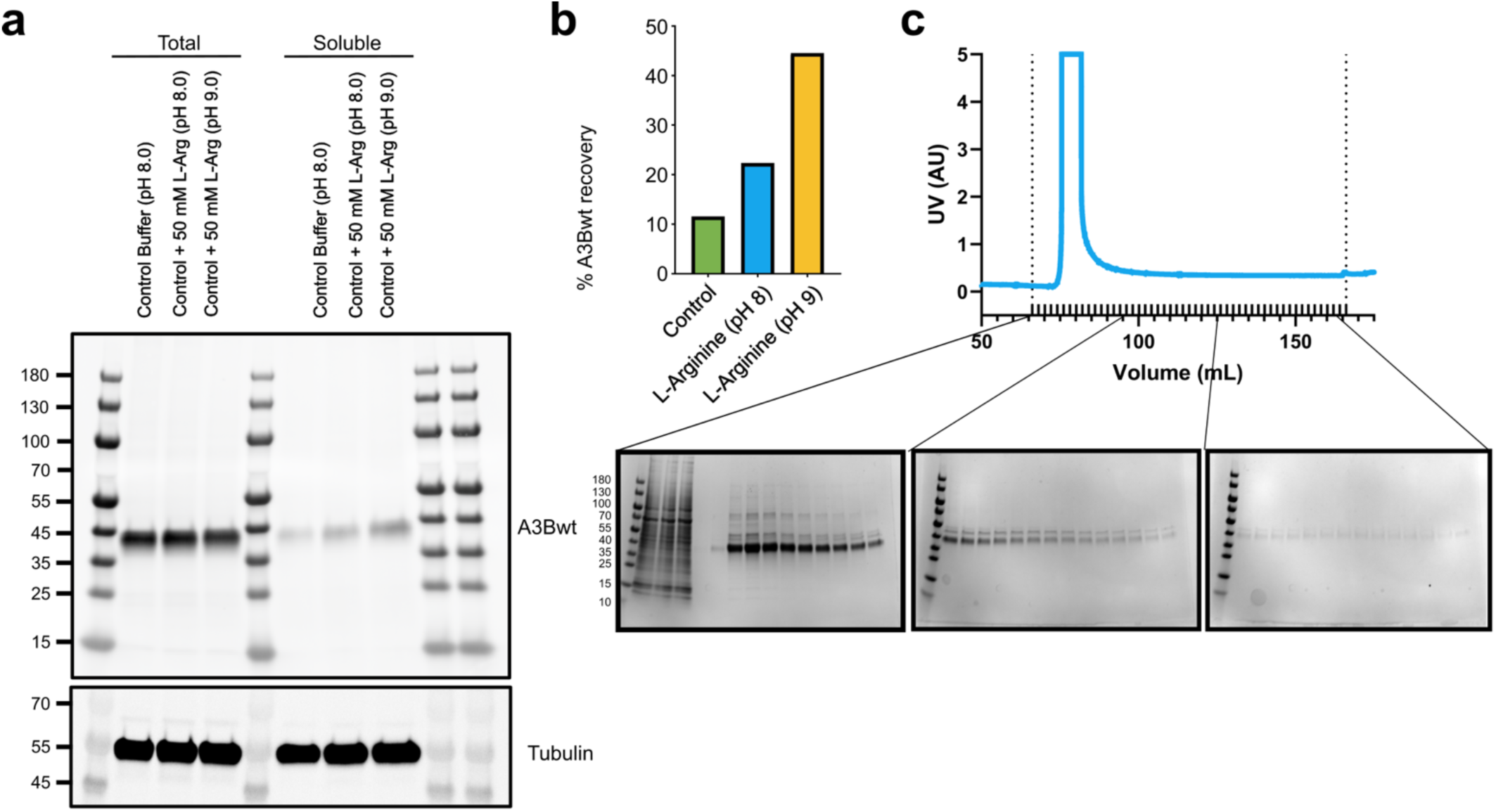
A3Bwt solubility testing and purification. **a**, Immunoblot detecting A3Bwt or tubulin in the total and clarified lysates in a control lysis buffer or the lysis buffer supplemented with 50 mM L-arginine at pH 8 or 9. **b**, Quantification of the immunoblot from panel a. Percent recovery was defined as the ratio of the integrated areas under the peaks of the soluble fraction to the total fraction. **c**, Top, Ni-NTA chromatogram of a representative purification of A3Bwt. Elution was carried out at a constant imidazole concentration of 400 mM. Bottom, SDS-PAGE gels of all fractions collected between the dotted lines in the chromatogram. A3Bwt is detectable by Coomassie stain over the entire elution.

**Extended Data Figure 2:**
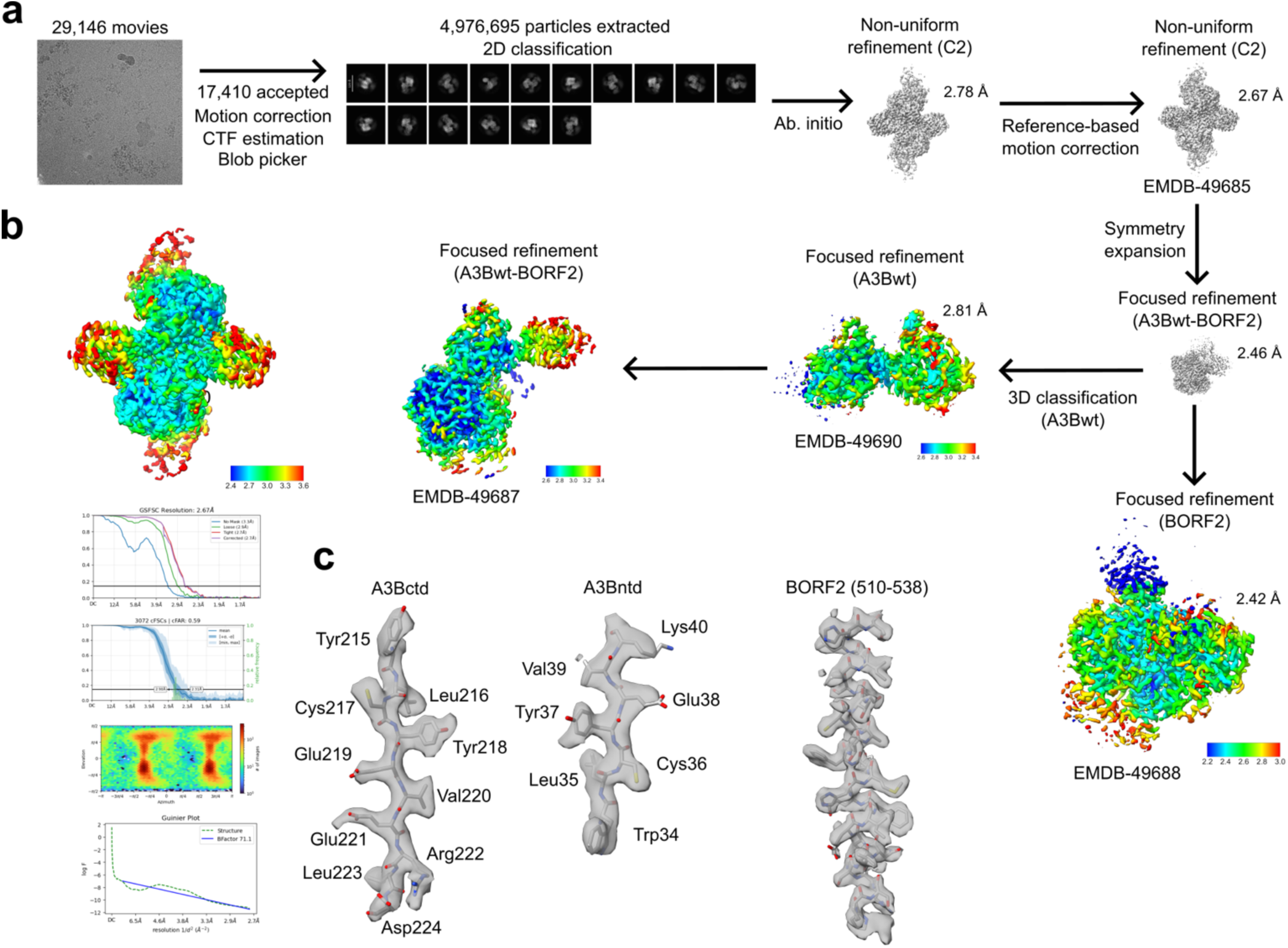
CryoEM data processing pipeline. **a**, Data processing pipeline for the A3Bwt-BORF2 dataset. **b**, Top, CryoEM map of the heterotetrameric A3Bwt-BORF2 complex refined with C2 symmetry applied and colored by local resolution. Bottom, GS-FSC plot, conical FSC plot, orientation distribution plot, and Guinier plot of this refinement. **c**, Model-map fits for representative β-strands (left, middle) or α-helices (right) from A3Bntd, A3Bctd, and BORF2, respectively.

**Extended Data Figure 3:**
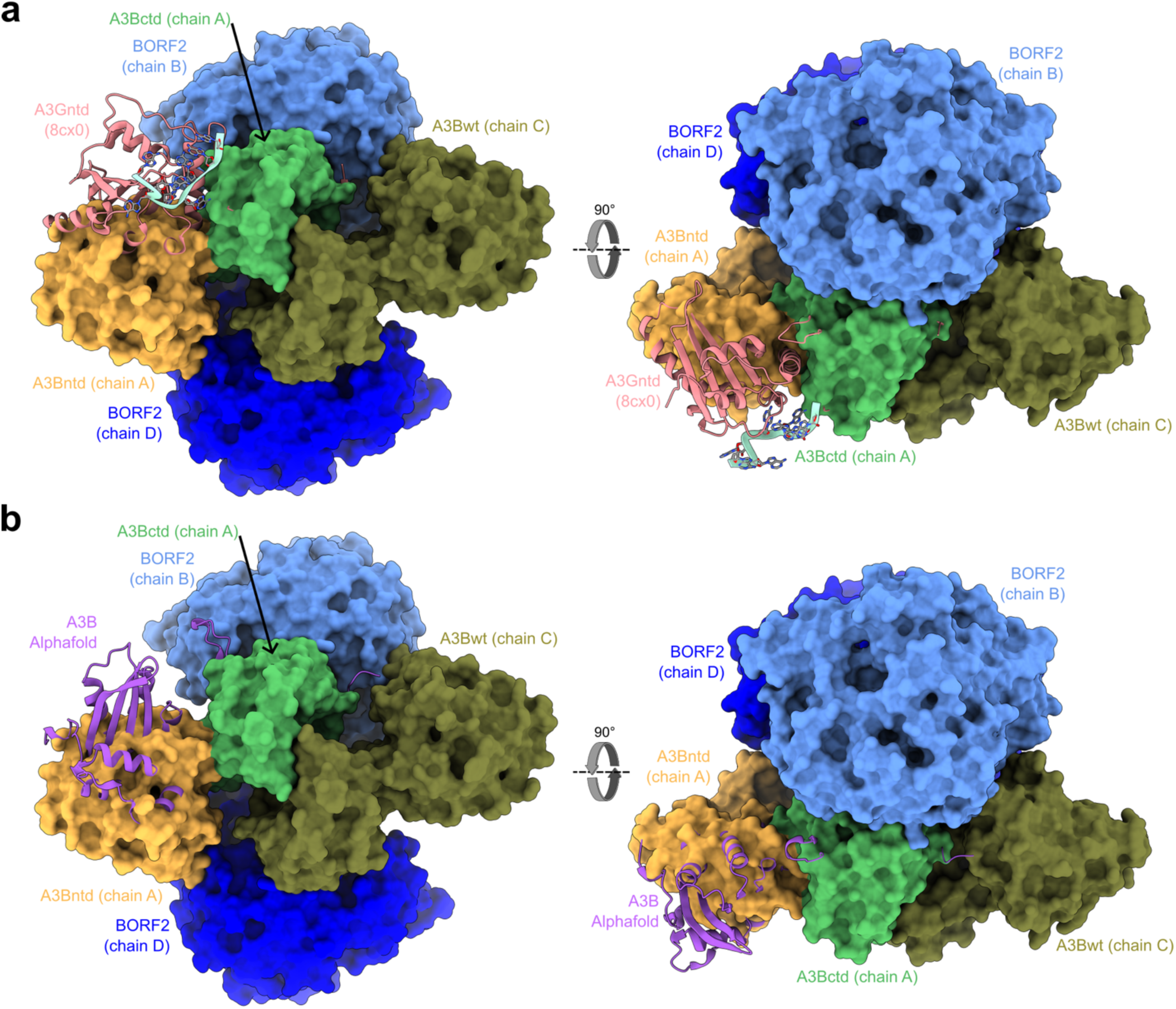
Analysis of A3G or AlphaFold-predicted A3B modeled onto the A3Bwt-BORF2 complex. **a**, Two views of the A3Bwt-BORF2 complex, colored as in Figure 3, rotated by 90°. A3G from the A3G-ssRNA-Vif complex (pdb: 8cx0, pink) was aligned to one copy of A3Bwt using the CTD. **b**, Two views of the A3Bwt-BORF2 complex, colored as in Figure 3, rotated by 90°. A3B-AF (purple) was aligned to one copy of A3Bwt using the CTD. Neither A3G nor A3B-AF shows a steric clash of NTD with the rest of the complex in these hypothetical models, suggesting that A3Bwt in these conformations could have been accommodated.

**Extended Data Figure 4:**
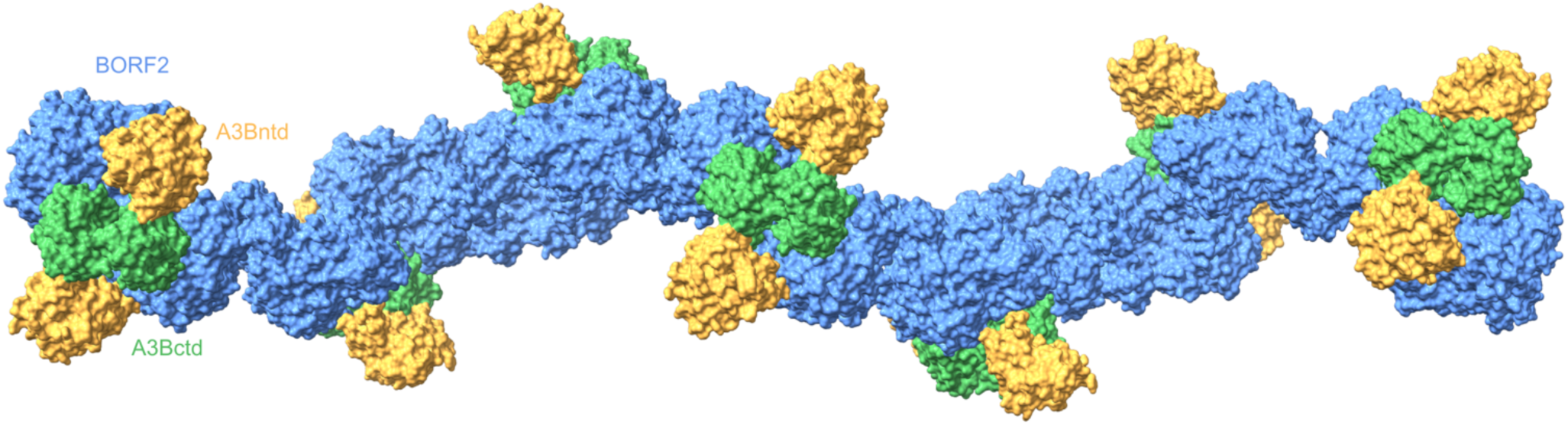
The A3Bwt-BORF2 “sponge” model A model of a BORF2 oligomer (blue) with A3Bwt (NTD = orange, CTD = green) bound, generated by using the noncanonical BORF2 dimer interface as observed in the A3Bctd-BORF2 structure (7rw6, Figure 2g) to link together multiple copies of the A3Bwt-BORF2 complex.

## References

1. Harris, R.S. & Dudley, J.P. APOBECs and virus restriction. Virology 479-480C, 131–145 (2015).

2. Lovsin, N., Gangupam, B. & Bergant Marusic, M. The intricate interplay between APOBEC3 proteins and DNA tumour viruses. Pathogens 13(2024).

3. Stavrou, S. & Ross, S.R. APOBEC3 proteins in viral immunity. J Immunol 195, 4565–70 (2015).

4. Silvas, T.V. & Schiffer, C.A. APOBEC3s: DNA-editing human cytidine deaminases. Protein Sci 28, 1552–1566 (2019).

5. Cheng, A.Z. et al. APOBECs and herpesviruses. Viruses 13(2021).

6. Conticello, S.G. The AID/APOBEC family of nucleic acid mutators. Genome Biol 9, 229 (2008).

7. Uriu, K., Kosugi, Y., Suzuki, N., Ito, J. & Sato, K. Elucidation of the complicated scenario of primate APOBEC3 gene evolution. J Virol 95(2021).

8. Shi, K. et al. Structural basis for targeted DNA cytosine deamination and mutagenesis by APOBEC3A and APOBEC3B. Nat Struct Mol Biol 24, 131–139 (2017).

9. Kouno, T. et al. Crystal structure of APOBEC3A bound to single-stranded DNA reveals structural basis for cytidine deamination and specificity. Nat Commun 8, 15024 (2017).

10. Harjes, S. et al. Structure-guided inhibition of the cancer DNA-mutating enzyme APOBEC3A. Nat Commun 14, 6382 (2023).

11. Stenglein, M.D., Matsuo, H. & Harris, R.S. Two regions within the amino-terminal half of APOBEC3G cooperate to determine cytoplasmic localization. J Virol 82, 9591–9 (2008).

12. Pak, V., Heidecker, G., Pathak, V.K. & Derse, D. The role of amino-terminal sequences in cellular localization and antiviral activity of APOBEC3B. J Virol 85, 8538–47 (2011).

13. Auerbach, A.A., et al. Ancestral APOBEC3B nuclear localization is maintained in humans and apes and altered in most other Old World primate species. mSphere 7, e0045122 (2022).

14. Salamango, D.J. et al. APOBEC3B nuclear localization requires two distinct N-terminal domain surfaces. J Mol Biol 430, 2695–2708 (2018).

15. Cheng, A.Z. et al. A conserved mechanism of APOBEC3 relocalization by herpesviral ribonucleotide reductase large subunits. J Virol 93(2019).

16. Cheng, A.Z. et al. Epstein-Barr virus BORF2 inhibits cellular APOBEC3B to preserve viral genome integrity. Nat Microbiol 4, 78–88 (2019).

17. Moraes, S.N. et al. Evidence linking APOBEC3B genesis and evolution of innate immune antagonism by gamma-herpesvirus ribonucleotide reductases. Elife 11(2022).

18. Shaban, N.M. et al. Cryo-EM structure of the EBV ribonucleotide reductase BORF2 and mechanism of APOBEC3B inhibition. Sci Adv 8, eabm2827 (2022).

19. Burns, M.B. et al. APOBEC3B is an enzymatic source of mutation in breast cancer. Nature 494, 366–70 (2013).

20. Burns, M.B., Temiz, N.A. & Harris, R.S. Evidence for APOBEC3B mutagenesis in multiple human cancers. Nat Genet 45, 977–83 (2013).

21. Law, E.K. et al. The DNA cytosine deaminase APOBEC3B promotes tamoxifen resistance in ER-positive breast cancer. Sci Adv 2, e1601737 (2016).

22. Gupta, A. et al. APOBEC3 mutagenesis drives therapy resistance in breast cancer. Nat Genet 57, 1452–1462 (2025).

23. Carpenter, M.A. et al. Mutational impact of APOBEC3A and APOBEC3B in a human cell line and comparisons to breast cancer. PLoS Genet 19, e1011043 (2023).

24. Petljak, M. et al. Mechanisms of APOBEC3 mutagenesis in human cancer cells. Nature 607, 799–807 (2022).

25. Schreurs, M.A.C. et al. APOBEC3B protein expression associates with poor prognosis for breast cancer patients with ER-positive disease. Breast Cancer Res 28, 7 (2025).

26. Alexandrov, L.B. et al. The repertoire of mutational signatures in human cancer. Nature 578, 94–101 (2020).

27. Roberts, S.A. et al. An APOBEC cytidine deaminase mutagenesis pattern is widespread in human cancers. Nat Genet 45, 970–6 (2013).

28. Durfee, C. et al. Tobacco smoke carcinogens exacerbate APOBEC mutagenesis and carcinogenesis. *bioRxiv* 10.1101/2025.01.18.633716 (2025).

29. McCann, J.L. et al. APOBEC3B regulates R-loops and promotes transcription-associated mutagenesis in cancer. Nat Genet 55, 1721–1734 (2023).

30. Xiao, X. et al. Structural determinants of APOBEC3B non-catalytic domain for molecular assembly and catalytic regulation. Nucleic Acids Res 45, 7494–7506 (2017).

31. Chelico, L. & Adolph, M.B. Purification of enzymatically active APOBEC proteins from an insect cell expression system. Methods Enzymol 713, 31–68 (2025).

32. Belica, C.A. et al. A real-time biochemical assay for quantitative analyses of APOBEC-catalyzed DNA deamination. J Biol Chem 300, 107410 (2024).

33. Belica, C.A. et al. RADD: A real-time FRET-based biochemical assay for DNA deaminase studies. Methods Enzymol 705, 311–345 (2024).

34. Greene, B.L. et al. Ribonucleotide reductases: structure, chemistry, and metabolism suggest new therapeutic targets. Annu Rev Biochem 89, 45–75 (2020).

35. Shi, K., Carpenter, M.A., Kurahashi, K., Harris, R.S. & Aihara, H. Crystal structure of the DNA deaminase APOBEC3B catalytic domain. J Biol Chem 290, 28120–30 (2015).

36. Shi, K. et al. Conformational switch regulates the DNA cytosine deaminase activity of human APOBEC3B. Sci Rep 7, 17415 (2017).

37. Chen, Y., Mullally, C.D., Stefanovska, B. & Harris, R.S. HAMMER: hairpin-based APOBEC3A-mediated mRNA editing reporter. Nucleic Acids Res 54(2026).

38. Rieffer, A.E., Chen, Y., Salamango, D.J., Moraes, S.N. & Harris, R.S. APOBEC reporter systems for evaluating diNucleotide editing levels. CRISPR J 6, 430–446 (2023).

39. Jumper, J. et al. Highly accurate protein structure prediction with AlphaFold. Nature 596, 583–589 (2021).

40. Varadi, M. et al. AlphaFold protein structure database in 2024: providing structure coverage for over 214 million protein sequences. Nucleic Acids Res 52, D368–D375 (2024).

41. Li, Y.L. et al. The structural basis for HIV-1 Vif antagonism of human APOBEC3G. Nature 615, 728–733 (2023).

42. Ito, F. et al. Structural basis for HIV-1 antagonism of host APOBEC3G via Cullin E3 ligase. Sci Adv 9, eade3168 (2023).

43. Yang, H., Kim, K., Li, S., Pacheco, J. & Chen, X.S. Structural basis of sequence-specific RNA recognition by the antiviral factor APOBEC3G. Nat Commun 13, 7498 (2022).

44. Arii, J. et al. A viral APOBEC3 antagonist distinguishes HHV-6A from HHV-6B. Nat Commun 17(2026).

45. Brown, W.L. et al. A rabbit monoclonal antibody against the antiviral and cancer genomic DNA mutating enzyme APOBEC3B. Antibodies (Basel) 8(2019).

46. Shi, K. et al. Structural basis for recognition of distinct deaminated DNA lesions by endonuclease Q. Proc Natl Acad Sci U S A 118(2021).

47. Punjani, A., Rubinstein, J.L., Fleet, D.J. & Brubaker, M.A. cryoSPARC: algorithms for rapid unsupervised cryo-EM structure determination. Nat Methods 14, 290–296 (2017).

48. Rubinstein, J.L. & Brubaker, M.A. Alignment of cryo-EM movies of individual particles by optimization of image translations. J Struct Biol 192, 188–95 (2015).

49. Zivanov, J., Nakane, T. & Scheres, S.H.W. A Bayesian approach to beam-induced motion correction in cryo-EM single-particle analysis. IUCrJ 6, 5–17 (2019).

50. Zivanov, J., Nakane, T. & Scheres, S.H.W. Estimation of high-order aberrations and anisotropic magnification from cryo-EM data sets in RELION-3.1. IUCrJ 7, 253–267 (2020).

51. Casanal, A., Lohkamp, B. & Emsley, P. Current developments in Coot for macromolecular model building of Electron Cryo-microscopy and Crystallographic Data. Protein Sci 29, 1069–1078 (2020).

52. Emsley, P., Lohkamp, B., Scott, W.G. & Cowtan, K. Features and development of Coot. Acta Crystallogr D Biol Crystallogr 66, 486–501 (2010).

53. Liebschner, D. et al. Macromolecular structure determination using X-rays, neutrons and electrons: recent developments in Phenix. Acta Crystallogr D Struct Biol 75, 861–877 (2019).

54. Pintilie, G.D., Zhang, J., Goddard, T.D., Chiu, W. & Gossard, D.C. Quantitative analysis of cryo-EM density map segmentation by watershed and scale-space filtering, and fitting of structures by alignment to regions. J Struct Biol 170, 427–38 (2010).

55. Pettersen, E.F. et al. UCSF Chimera--a visualization system for exploratory research and analysis. J Comput Chem 25, 1605–12 (2004).

56. Zhang, K., Pintilie, G.D., Li, S., Schmid, M.F. & Chiu, W. Resolving individual atoms of protein complex by cryo-electron microscopy. Cell Res 30, 1136-1139 (2020).

57. Meng, E.C. et al. UCSF ChimeraX: Tools for structure building and analysis. Protein Sci 32, e4792 (2023).

58. Borges, L.B., Pereira, A.K.F., Silva, W.B.D., Monteiro, F.O.B. & Coutinho, L.N. Abdominal ultrasound in Saguinus ursulus. J Med Primatol 49, 307–314 (2020).

59. Goddard, T.D. et al. UCSF ChimeraX: Meeting modern challenges in visualization and analysis. Protein Sci 27, 14–25 (2018).

60. Bursch, K.L., Olp, M.D. & Smith, B.C. Analysis of continuous enzyme kinetic data using ICEKAT. Methods Enzymol 690, 109–129 (2023).

61. Brignole, E.J. et al. 3.3-A resolution cryo-EM structure of human ribonucleotide reductase with substrate and allosteric regulators bound. Elife 7(2018).

62. Westmoreland, D.E. et al. 2.6-A resolution cryo-EM structure of a class Ia ribonucleotide reductase trapped with mechanism-based inhibitor N(3)CDP. Proc Natl Acad Sci U S A 121, e2417157121 (2024).

